# Human genetic variants disrupt RGS14 nuclear shuttling and regulation of LTP in hippocampal neurons

**DOI:** 10.1101/2020.09.10.289991

**Authors:** Katherine E Squires, Kyle J Gerber, Matthew C Tillman, Daniel J Lustberg, Carolina Montañez-Miranda, Meilan Zhao, Suneela Ramineni, Christopher D Scharer, Feng-jue Shu, Jason P Schroeder, Eric A Ortlund, David Weinshenker, Serena M Dudek, John R Hepler

## Abstract

The human genome contains vast genetic diversity in the form of naturally occurring coding variants, yet the impact of these variants on protein function and physiology is poorly understood. RGS14 is a multifunctional signaling protein that suppresses synaptic plasticity in dendritic spines of hippocampal neurons. RGS14 also is a nucleocytoplasmic shuttling protein, suggesting that balanced nuclear import/export and dendritic spine localization are essential for RGS14 functions. We identified genetic variants L505R (LR) and R507Q (RQ) located within the nuclear export sequence (NES) of human *RGS14*. Here we report that RGS14 carrying LR or RQ profoundly impacts protein functions in hippocampal neurons and brain. Following nuclear import, RGS14 nuclear export is regulated by Exportin 1 (XPO1/CRM1). Remarkably, LR and RQ variants disrupt RGS14 binding to Gαi1-GDP and XPO1, nucleocytoplasmic equilibrium, and capacity to inhibit LTP. Variant LR accumulates irreversibly in the nucleus, preventing RGS14 binding to G proteins, localization to dendritic spines, and inhibitory actions on LTP induction, while variant RQ exhibits a mixed phenotype. When introduced into mice by CRISPR/Cas9, RGS14-LR protein expression was detected predominantly in the nuclei of neurons within hippocampus, central amygdala, piriform cortex, and striatum, brain regions associated with learning and synaptic plasticity. Whereas mice completely lacking RGS14 exhibit enhanced spatial learning, mice carrying variant LR exhibit normal spatial learning, suggesting that RGS14 may have distinct functions in the nucleus independent from those in dendrites and spines. These findings show that naturally occurring genetic variants can profoundly alter normal protein function, impacting physiology in unexpected ways.

## Introduction

Despite 99.9% similarity across all human genomes, the human population nevertheless exhibits vast genetic diversity responsible for disease predisposition and distinct human traits. This genetic diversity is not captured by canonical reference protein sequences and, despite selective pressure, missense variations within exonic coding regions frequently arise in modular protein domains that can have profound impact on protein functions. While many studies focus on monogenetic variations in the context of severe diseases and phenotypes, rare variants may actually play a more important role in susceptibility, particularly for complex diseases (1-8). Many of these rare variants are predicted to alter protein function which could impact linked physiology. Thus, exploration of rare missense human variants in cell and animal models can serve as a useful tool to reveal novel protein functions relevant to human physiology.

The *<u>R</u>*egulators of *<u>G</u>* protein *<u>S</u>*ignaling (RGS) proteins modulate GPCR/G protein signaling (9-11) which controls many aspects of neurobiology and cognition (12), including synaptic plasticity and learning (13,14). GPCR signaling is tightly regulated by RGS proteins which catalyze the off-rate of active Gα-GTP, and thus RGS proteins also control many aspects of synaptic plasticity (15) in both physiology and disease states (16). All RGS proteins (20 classical family members) share a conserved ∼120 amino acid RGS domain, and many contain accessory domains which act to regulate different aspects of cellular function (9,10), such as MAP kinase signaling (17-19). Although canonical reference sequences are used to define and study RGS and other proteins, our recent report identified considerable genetic diversity within each human RGS protein family member, with many encoding missense variants (4). These missense variants can have potentially deleterious effects on RGS protein function with physiological consequences (4).

In our report (4), RGS14 emerged as one family member with considerable genetic variance. RGS14 is a dynamically regulated multifunctional signaling protein that naturally suppresses synaptic plasticity in hippocampal neurons (20,21). In addition to the canonical RGS domain, RGS14 also contains other signaling domains including two tandem Ras/Rap binding domains (R1 and R2) which bind active H-Ras-GTP and Rap2A-GTP to regulate MAP kinase signaling (17-19). RGS14 also contains a *<u>G p</u>*rotein *<u>r</u>*egulatory (GPR) motif, which governs RGS14 subcellular localization by interactions with inactive Gαi1/3-GDP at the plasma membrane (22,23). The GPR motif recruits RGS14 to the plasma membrane where RGS14 can exert its actions on downstream effectors (24,25). Whereas the canonical GAP function of RGS14 towards Gαi/o-GTP is dependent on the RGS domain, RGS14 subcellular localization and distribution is dependent on the GPR motif (25). Besides engaging G protein signaling at the plasma membrane, RGS14 is also a nucleocytoplasmic shuttling protein whose subcellular movement is regulated by a nuclear localization sequence (NLS) and a nuclear export sequence (NES) (25,26). The NES is encoded within the GRP motif and dictates RGS14 nuclear export (25,27). Thus, the GPR motif, and the NES within, are critically important for RGS14 subcellular localization, distribution, and cellular functions.

We previously reported that, in mouse brain, RGS14 is highly expressed within the small enigmatic CA2 subregion of the hippocampus, but not in the neighboring CA1 or CA3 subregions (20,28). While CA1 is known for its robust long-term potentiation (LTP), a form of synaptic plasticity that is thought to underlie several forms of learning and memory (29), canonical LTP is absent from CA2 neurons where RGS14 is naturally expressed (30). Of note, RGS14 localizes predominately to spines and dendrites in CA2 neurons, making it well positioned to modulate LTP and synaptic plasticity. In support of this idea, we reported that genetic ablation of RGS14 resulted in robust LTP in CA2 (mirroring that of CA1), with no changes in CA1, and that introduction of RGS14 into CA1 neurons blocked LTP there (20,21). Further, loss of RGS14 conferred an enhancement in the rate of spatial learning, a hippocampal-dependent task (20), demonstrating that RGS14 is a natural suppressor of LTP linked to spatial learning. However, whether these findings extended to primates, specifically humans, remained an open question.

Our recent report detailed the regional, cellular and subcellular localization of RGS14 in the primate brain (31). We observed that RGS14 is expressed in hippocampus, as expected, but also in amygdala and striatum (28). We also observed for the first-time native expression of RGS14 in the nuclei of a subpopulation of striatal neurons in primates. This observation of native nuclear RGS14 is consistent with earlier reports demonstrating that RGS14 is a dynamically regulated nucleocytoplasmic shuttling protein (25,26). Of note, we also observed possible RGS14 splice variants lacking the GPR/NES, which would be expected to target this form of RGS14 to the nucleus. The discovery of a subpopulation of nuclear RGS14 in monkey brain raised numerous questions including: 1) how is RGS14 nuclear localization controlled, 2) what is the impact of nuclear localization on RGS14 cellular functions, and 3) if putative splice variants exist in monkey which shift the nucleocytoplasmic balance, do naturally occurring rare human variants shift nucleocytoplasmic shuttling as well?

To explore these questions, we took advantage of human genetic variant data collected from over 130,000 individuals to examine the contribution of naturally occurring mutations on RGS14 functions (32,33). Human RGS14 contains many genetic variants observed throughout the gene. Here we report that two human genetic variants, specifically L505R (LR) and R507Q (RQ) located within the NES of RGS14, profoundly impact RGS14 functions in mouse hippocampal neurons and brain. We provide evidence that the GPR motif of RGS14, and the NES encoded within, are critically important for RGS14 subcellular localization and capacity to prevent LTP induction. Variants LR and RQ disrupt RGS14 binding to Gαi1-GDP and XPO1, nucleocytoplasmic equilibrium, and capacity to inhibit LTP in hippocampal neurons. We introduced RGS14 variant LR into mice by CRISPR/Cas9 and found this variant to direct RGS14 protein to the nuclei of neurons in the hippocampus, central amygdala, and striatum, brain regions rich in synaptic plasticity. Whereas mice lacking RGS14 are reported to exhibit enhanced spatial learning (20), those expressing variant LR exhibit normal spatial learning, suggesting that RGS14 has distinct functions in the nucleus versus dendrites and postsynaptic dendritic spines. These findings demonstrate that naturally occurring genetic variants can alter normal protein functions in unexpected ways, and in the case of RGS14, with profound physiological consequences in the hippocampus and likely other brain regions.

## Results

### The GPR motif is critical for RGS14 capacity to inhibit synaptic plasticity in hippocampal neurons

We previously reported that RGS14 is selectively expressed in neurons of hippocampal area CA2 where it acts as a natural suppressor of synaptic plasticity in mouse brain (20,21). Unlike area CA1, which exhibits robust LTP and low levels of RGS14, CA2 pyramidal neurons fail to exhibit LTP following Schaffer collateral stimulation (30). Loss of RGS14 (RGS14 KO) restores LTP in CA2 neurons in acute mouse hippocampal slices (20,21), and exogenous introduction of RGS14 into area CA1 completely blocks synaptic plasticity there (21), suggesting that RGS14 can engage postsynaptic signaling pathways shared by both CA1 and CA2 neurons to inhibit synaptic plasticity. Based on these observations, we sought to isolate which structural region(s) of RGS14 is/are most important for regulating LTP in hippocampal neurons (Figure 1).

RGS14 is a multifunctional protein containing: 1) an RGS domain which binds active Gαi/o-GTP; 2) tandem Ras-binding domains (R1 and R2) which bind active H-Ras-GTP and Rap2A-GTP; and 3) a GPR (G) motif which binds inactive Gαi1/3-GDP (17-19,22-25) (Figure 1A). We initiated studies to determine which domain(s) and protein interaction(s) are most critical for RGS14 regulation of LTP. Because RGS14 blocks LTP in CA1 neurons when introduced by ectopic expression (21), we used this system to examine the effects of targeted loss-of-function mutations (Figure S1) of RGS14 on LTP in CA1 neurons. Each of these mutations allowed for specific RGS14 functions to be blocked while leaving others intact. We generated adeno-associated viruses (AAV) expressing YFP and wild type YFP-RGS14 as negative and positive controls (Figure S1). We also generated AAV expressing: 1) YFP-RGS14 E92A/N93A (EN/AA) in the RGS domain, which blocks Gαi/o-GTP binding (34); 2) YFP-RGS14-R333L in the R1 domain, which blocks H-Ras-GTP binding (18); and 3) YFP-RGS14-Q515A/R516A (QR/AA) in the GPR motif, which blocks Gαi1/3-GDP binding (24,34) (Figure S1 and Figure 1A). Virus was expressed in CA1 of hippocampal slice cultures (Figure 1B-C) and, excitatory postsynaptic currents (EPSCs) were measured by whole cell voltage clamp recordings as performed previously on acute slices (20,21,30). While control and YFP-only recordings show robust LTP in response to a pairing protocol, expression of wild type YFP-RGS14 fully blocks LTP when ectopically expressed in CA1 hippocampal neurons (Figure 1D-F) as expected (21), demonstrating that RGS14 is sufficient to suppress LTP. Inactivation of the RGS domain or R1 domain did not alter RGS14 capacity to suppress LTP (Figure 1G-H), suggesting these domains and linked protein interactions are less critical for this RGS14 function. However, inactivation of the GPR (G) motif was able to block RGS14 function and allow LTP in CA1 neurons (Figure 1I). These results indicate that the GPR motif, a regulator of RGS14 localization within the cell (24,34), is critical for RGS14 suppression of synaptic plasticity, prompting the notion that spatial dynamics are a critical mediator of RGS14 function.

**Figure 1.**
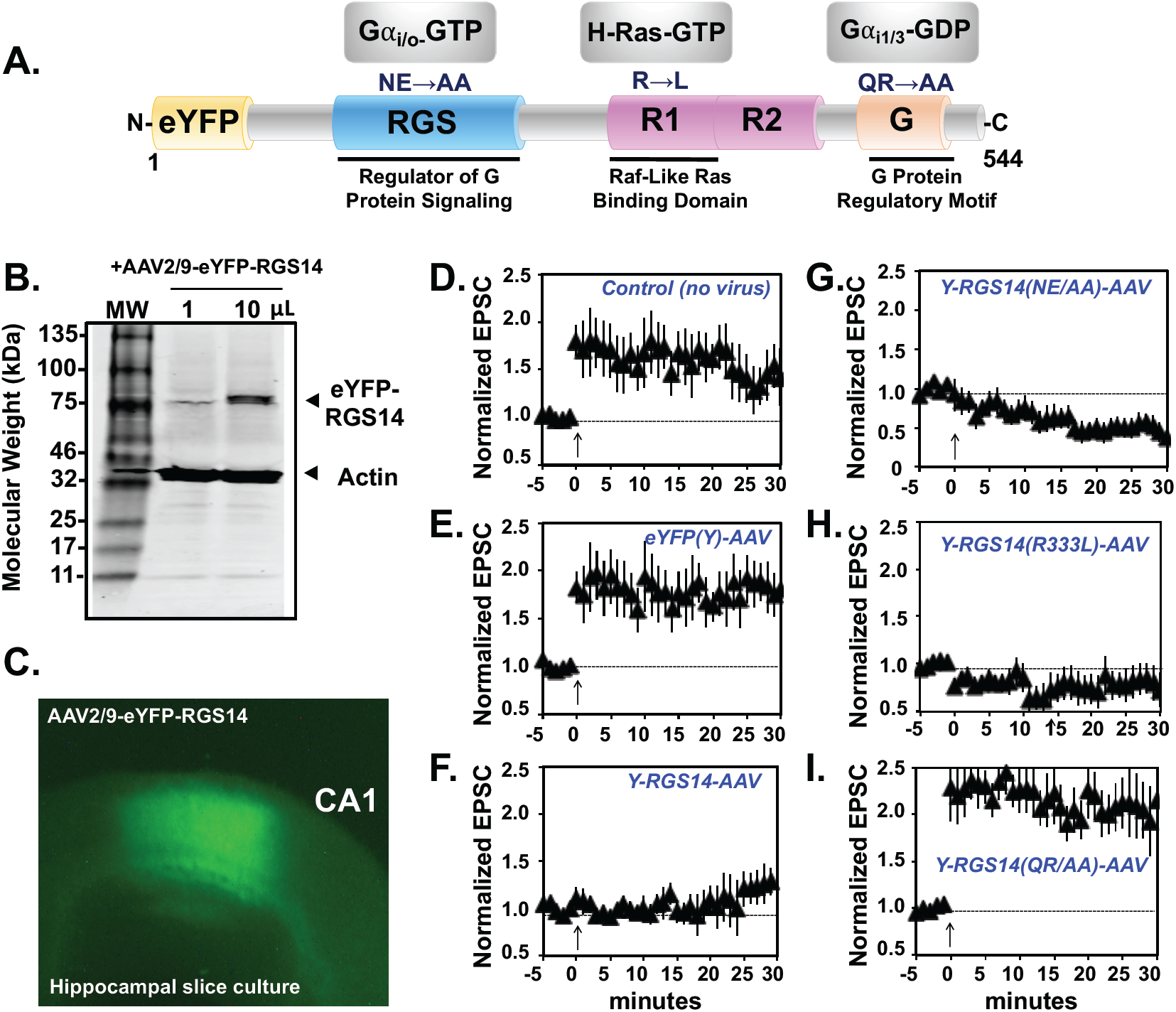
The GPR motif is critical for RGS14 suppression of LTP. **A)** RGS14 is a multifunctional signaling protein which contains an RGS domain that binds active Gαi/o-GTP, an R1 domain that binds H-Ras-GTP, and the GPR motif (G) that binds inactive Gαi1/3-GDP. Amino acid substitutions are indicated for each domain which confer a loss-of-function phenotype for that specific domain (see Figure S1). **B)** eYFP-RGS14 constructs, wild type and mutant, were placed in AAV and, following infection, expressed as full-length intact proteins in hippocampal neurons, as detected by coomassie blue stain. **C)** eYFP-RGS14 is expressed in area CA1 of cultured hippocampal slices following infection with AAV. **D-F)** Long term potentiation (LTP) was performed by pairing Schaffer collateral input with CA1 neuron depolarization (time indicated with the arrow) in cultured hippocampal slices expressing AAV-eYFP-RGS14 constructs. Following the LTP-pairing protocol, EPSCs were assessed for potentiation. Control CA1 neurons (n =12) and those expressing YFP alone (n=6) showed robust LTP, while eYFP-RGS14 (n=5) suppressed induction of LTP. **G-I)** Although the eYFP-RGS14 NE/AA (RGS domain knockout; n = 6) and eYFP-RGS14 R/L (R1 domain knockout; n=5) suppressed LTP similar to WT RGS14, the eYFP-RGS14 GPR motif knockout QR/AA (n=6) failed to suppress LTP, suggesting the GPR motif is a critical mediator of RGS14 function. Data is presented as mean +/-SD.

### Naturally occurring rare human genetic variants within the GPR motif disrupt RGS14 interactions with Gαi1-GDP

The mutations used above to disrupt RGS14 GPR motif function were rationally designed within the context of the canonical RGS14 sequence. Based on the newly appreciated importance of the GRP motif for RGS14 function in LTP (Figure 1), we next explored whether naturally occurring human variants existed within the GPR motif that might impact RGS14 protein function. We recently reported the existence of numerous missense variants in the human RGS14 sequence, including many within the GPR motif (4) that are predicted to disrupt function. For this, we accessed the Genome Aggregation Database (GnomAD, version 2.0) for all RGS14 missense (amino acid change) and silent (DNA change but no amino acid change) variants. We observed a relatively even distribution of missense and silent variants throughout the RGS14 gene (Figure 2A, missense on top, silent on bottom). Germane to RGS14 regulation of G protein signaling, multiple naturally occurring rare variants were found within the GPR motif (Figure 2A, GPR motif and sequence expanded with variants shown on top in red).

**Figure 2.**
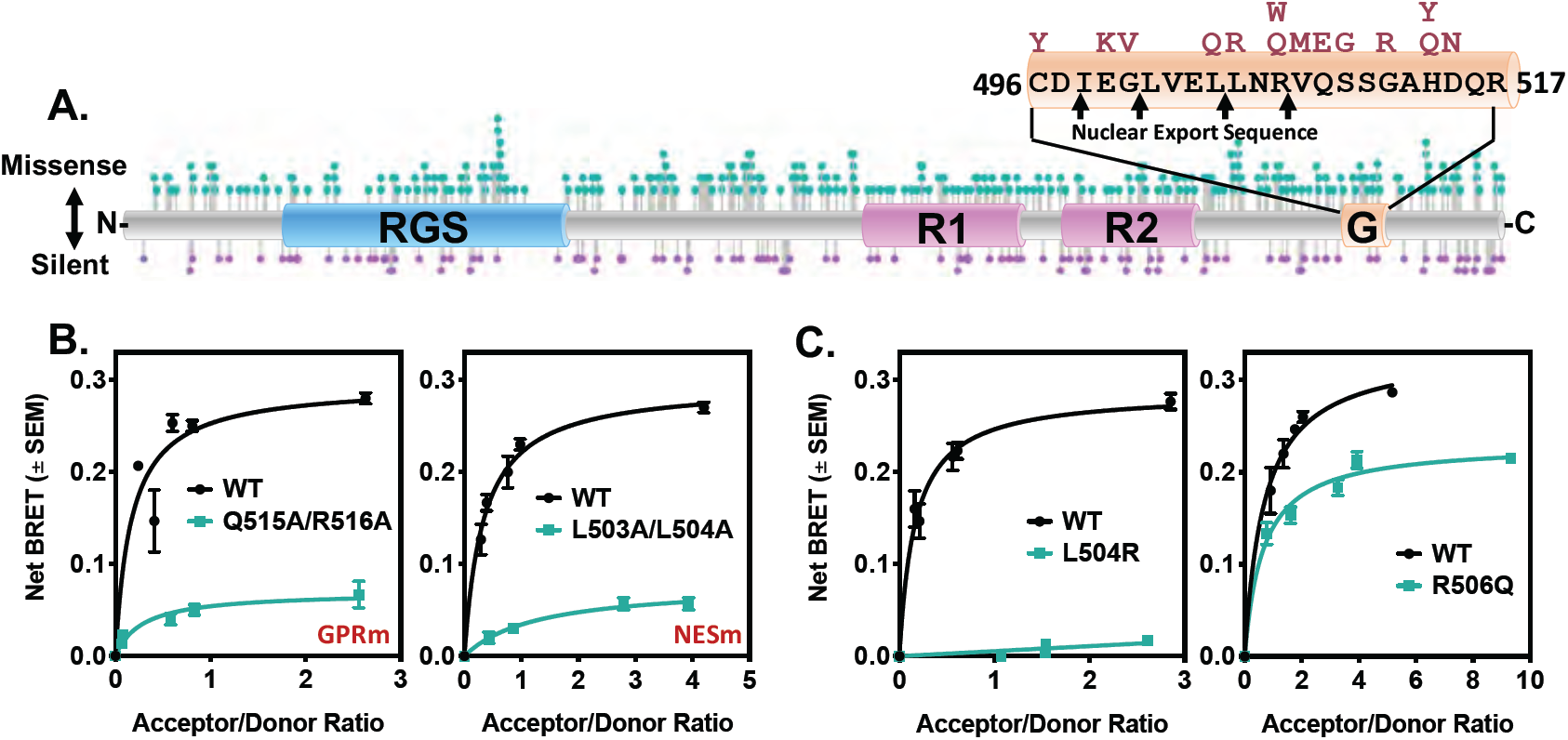
Rare human genetic variants within the GPR and NES motifs of RGS14 disrupt its association with Gαi1-GDP, suggestive of aberrant cellular trafficking. **A)** RGS14 missense and silent variants were obtained from GnomAD (Broad Institute) and plotted onto the sequence of RGS14 using Lollipops (https://github.com/pbnjay/lollipops) as described previously (4). Missense variants are plotted on top in teal, with silent variants on the bottom in purple. The GPR motif, which contains an embedded NES, has human variants as shown (red). **B-C)** BRET measurement of RGS14-Luc interaction with Gαi1-eYFP-GDP. RGS14-Luc is recruited from the cytoplasm to the plasma membrane by inactive Gαi1-eYFP-GDP (24). Inactivation of the GPR in RGS14-Luc (Q515A/R516A; GPRm; n=3) prevents Gαi1-eYFP-GDP binding and recruitment to the plasma membrane. Similarly, inactivation of the NES (L503A/L504A; NESm; n=3) prevents RGS14 from being accessible for recruitment in the cytoplasm, and therefore cannot associate with Gαi1-GDP at the plasma membrane. Variants L504R or R506Q were placed in RGS14-Luc and tested for their capacity to associate with Gαi1-eYFP-GDP by BRET, indicative of proper recruitment to the plasma membrane. L504R (n=3) completely abolished Gαi1-GDP association, while R506Q (n=3) exhibited a partial reduction in Gαi1-GDP association. Data is presented as mean +/-SEM.

The GPR motif binds inactive Gαi1-GDP and serves to recruit and stabilize RGS14 at the plasma membrane (18,24). RGS14:Gαi1-GDP interactions are readily detectable by BRET and are reflective of functional RGS14 trafficking between the cytosol and the plasma membrane (24). Wild type RGS14-Luc binds Gαi1-YFP and saturates the net BRET signal in a concentration-dependent manner (Figure 2B, black line). Importantly, the GPR motif has embedded within it a nuclear export sequence (NES, Figure 2A), which shuttles nuclear RGS14 to the cytoplasm (26). The NES is therefore also an important mediator of RGS14 subcellular localization, as it provides a cytoplasmic pool of RGS14 for Gαi1-GDP recruitment and co-localization at the plasma membrane. When the GPR motif is mutated (Q515A/R516A) such that it cannot bind Gαi1-GDP (GPRm) (24,35) (Figure S1) or when the NES is mutated (L503A/L504A) such that RGS14 is incapable of nuclear export (NESm) (25,26), RGS14-Luc ceases to bind Gαi1-YFP as measured by BRET, indicating aberrant subcellular trafficking (Figure 2B, teal lines). Together, these findings confirm that this screening assay is a sensitive measure of Gαi1-GDP binding and RGS14 subcellular spatial dynamics.

We identified 13 human variants in the RGS14 GPR motif (Figure 2A). Each of these variants were measured for their interactions with Gαi1-GDP by BRET and we found that, while the majority did not affect the Gαi1-YFP BRET signal (Figure S2), several did. Two rare variants (defined here as <2% frequency) emerged as especially interesting based on their effect and population frequency. Human RGS14 variant L505R (L504R in rat sequence), which is very rare (found in 0.006% of East Asian population [GnomAD v2.0]), completely abolished interaction with Gαi1-YFP (Figure 2C). Human RGS14 variant R507Q (R506Q in rat sequence; found in 1.25% of Ashkenazi Jewish population, among other populations [GnomAD v2.0]) has a submaximal reduction in Gαi1-YFP interaction and BRET signal.

### RGS14 is a nuclear shuttling protein in primary hippocampal neurons

We and others have reported that recombinant RGS14 shuttles into and out of the nucleus in cultured cell lines (25-27) and neurons (36). As noted, RGS14 contains a functional nuclear localization sequence (NLS) between the RGS domain and the first Ras-binding domain (R1), and a functional NES embedded within the GPR motif of RGS14 (25,26) (Figure 3A). The GPR motif and NES play an important role in determining the subcellular localization and spatial dynamics of RGS14 when transfected into HEK cells (18,25). However, the dynamic subcellular behavior of RGS14 in neurons has not been well described. Of note, we reported that RGS14 exists *natively* in the nuclei of striatal neurons in primate brain (31). In this same study, we also noted the existence of a possible splice variant lacking the NES, suggesting a pool of RGS14 could reside in the nuclei of these neurons. Based on this information, we next sought to determine the subcellular localization and nucleocytoplasmic shuttling behavior of RGS14 within primary hippocampal neurons.

**Figure 3.**
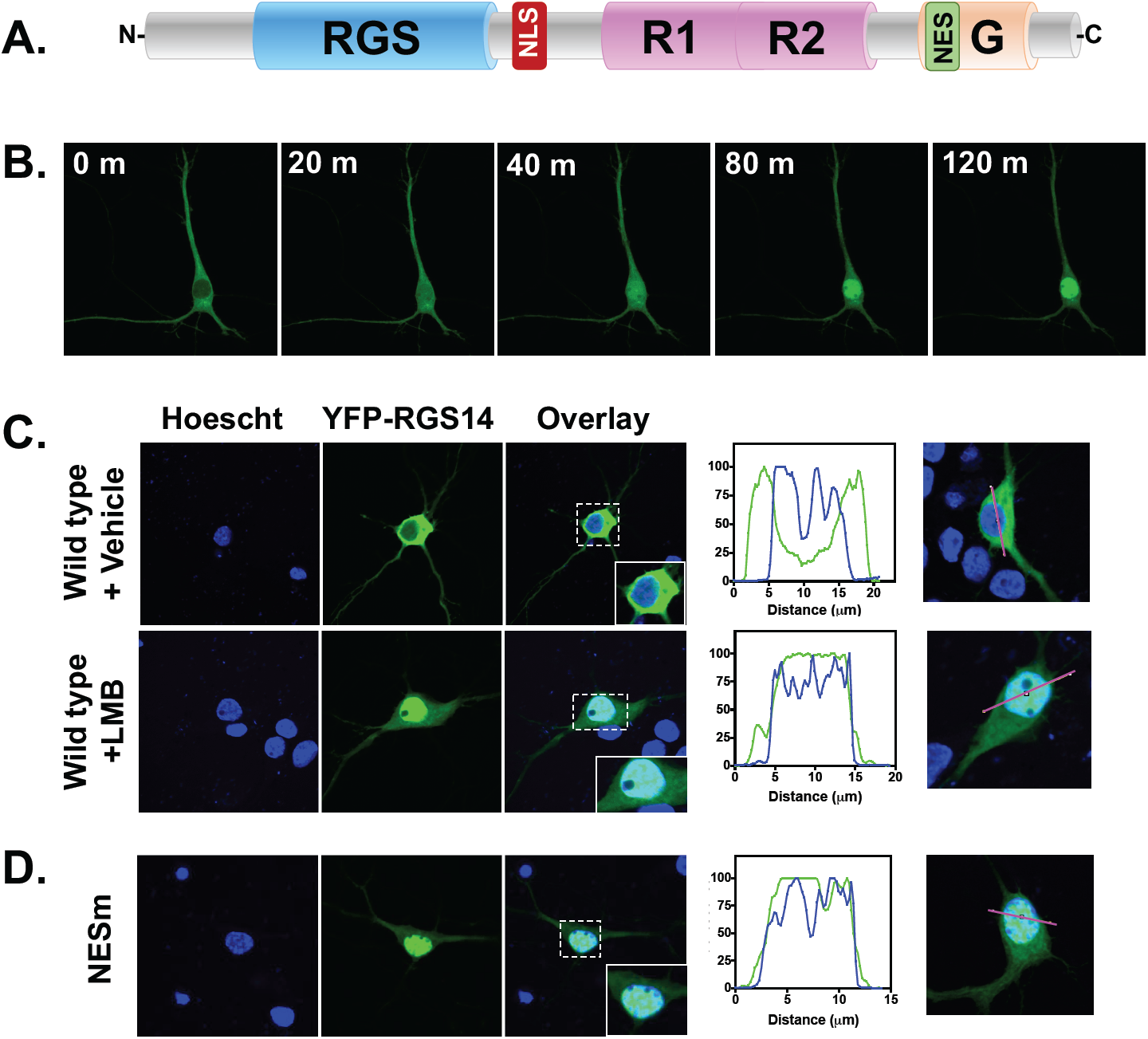
RGS14 is a nucleocytoplasmic shuttling protein in hippocampal neurons. **A)** RGS14 rapidly shuttles into and out of the nucleus in dividing cells directed by a nuclear localization sequence (NLS) and nuclear export sequence (NES). Studies validated that RGS14 followed these same cellular dynamics in primary hippocampal neurons. **B)** Neurons were infected with AAV-eYFP-RGS14 for 18 hours and then imaged using live cell confocal microscopy. Leptomycin B (LMB) was added (final concentration 20 nM) and YFP signal was measured. At time zero, YFP-RGS14 is entirely cytosolic. After 20 minutes, YFP-RGS14 was detectable in the nucleus, at 40 minutes YFP-RSG14 was mostly nuclear, and by 80-120 minutes YFP-RGS14 was entirely nuclear. **C)** Wild type YFP-RGS14 does not colocalize with Hoechst in vehicle treated conditions, but colocalizes with Hoechst entirely under LMB conditions. **D)** Mutation of the NES (L503A/L504A, NESm) in RGS14 sequesters RGS14 entirely in the nucleus under vehicle treated conditions. Approximately 10 images were collected per condition and representative images are shown.

Neurons (DIV 8) were transfected with AAV-hSyn-YFP-RGS14 for 18 hours and treated with the nuclear export inhibitor Leptomycin B (LMB) over two and a half hours, concurrent with imaging by live cell confocal microscopy. We found that cytoplasmic RGS14 translocated to the nucleus within 40 minutes and was maximally nuclear by two hours (Figure 3B), suggesting that cytoplasmic pools of RGS14 are constantly targeted for nuclear import/export with an equilibrium favoring the cytosol. These results are consistent with our recently published report demonstrating that nuclear import of RGS14 is regulated by 14-3-3γ in neurons (36).

We next verified co-localization of RGS14 with a nuclear marker, Hoescht, as well as validated the functional status of the RGS14 NES motif in neurons. DIV 8 neurons were again transfected with AAV-YFP-RGS14 WT or the NES mutant (NESm) incapable of being shuttled out of the nucleus (25,26). Neurons were treated with vehicle or LMB for two hours, then fixed and stained with Hoechst, and YFP signal was assessed by confocal microscopy. Vehicle-treated neurons exhibited robust YFP-RGS14 expression that filled the soma, dendrites, and spines of the neurons, but did not overlap with the Hoechst-stained nucleus. In contrast, YFP-RGS14 WT colocalized entirely with nuclear Hoechst in LMB-treated neurons (Figure 3C). Immunoblot analysis verified that full-length YFP-RGS14 was expressed (Figure S3), confirming this nuclear localization was not an effect of post-translationally modified (e.g., truncated) RGS14. LMB strongly and specifically inhibits nuclear export receptor Exportin 1 (XPO1, also known as CRM1) (37). The XPO1-binding motif is a leucine-rich sequence (38). We and others previously showed that mutating two leucines (L503A/L504A) as mentioned above (Figure 2B) in the RGS14 GPR motif (LL/AA) inhibited nuclear export of RGS14 in immortal cells (25,26). Thus, we examined in complimentary experiments whether RGS14 nuclear shuttling in primary hippocampal neurons occurred through the same mechanism. AAV-hSyn-YFP-RGS14 LL/AA (NESm) was transfected into DIV 8 neurons and stained with Hoechst as before. Again, we verified expression of a single band corresponding to full-length YFP-RGS14 NESm by immunoblot (Figure S3). We found that under vehicle-treated conditions, YFP-RGS14 NESm sequestered in the nuclei and colocalized with Hoechst, indistinguishable from the LMB-treated WT RGS14 (Figure 3D). These findings confirm that RGS14 nucleocytoplasmic shuttling behaves similarly in primary hippocampal neurons as it does in other cells, and that RGS14 nuclear export is mediated by XPO1.

### Human variants LR and RQ mislocalize RGS14 to the nucleus in primary hippocampal neurons

As illustrated in Figure 2, RGS14 genetic variants LR and RQ exhibit reduced Gαi1-YFP binding. Based on this observation, we hypothesized that RGS14 variants might also exhibit aberrant cellular trafficking, either via reduced plasma membrane recruitment or reduced nuclear shuttling. To distinguish between these possibilities, we transfected DIV 8 neurons with AAV-hSyn-YFP-RGS14 WT, AAV-hSyn-YFP-RGS14 RQ, or AAV-hSyn-YFP-RGS14 LR for 18 hours, then fixed and stained them with Hoechst as before. Confocal imaging captured the subcellular localization of wild type or variant RGS14 within the neuron. Wild type RGS14 localized to the soma, dendrites, and spines. In stark contrast, RGS14 LR concentrated within the nucleus while the RQ variant exhibited a mixed phenotype (Figure 4A-B).

**Figure 4.**
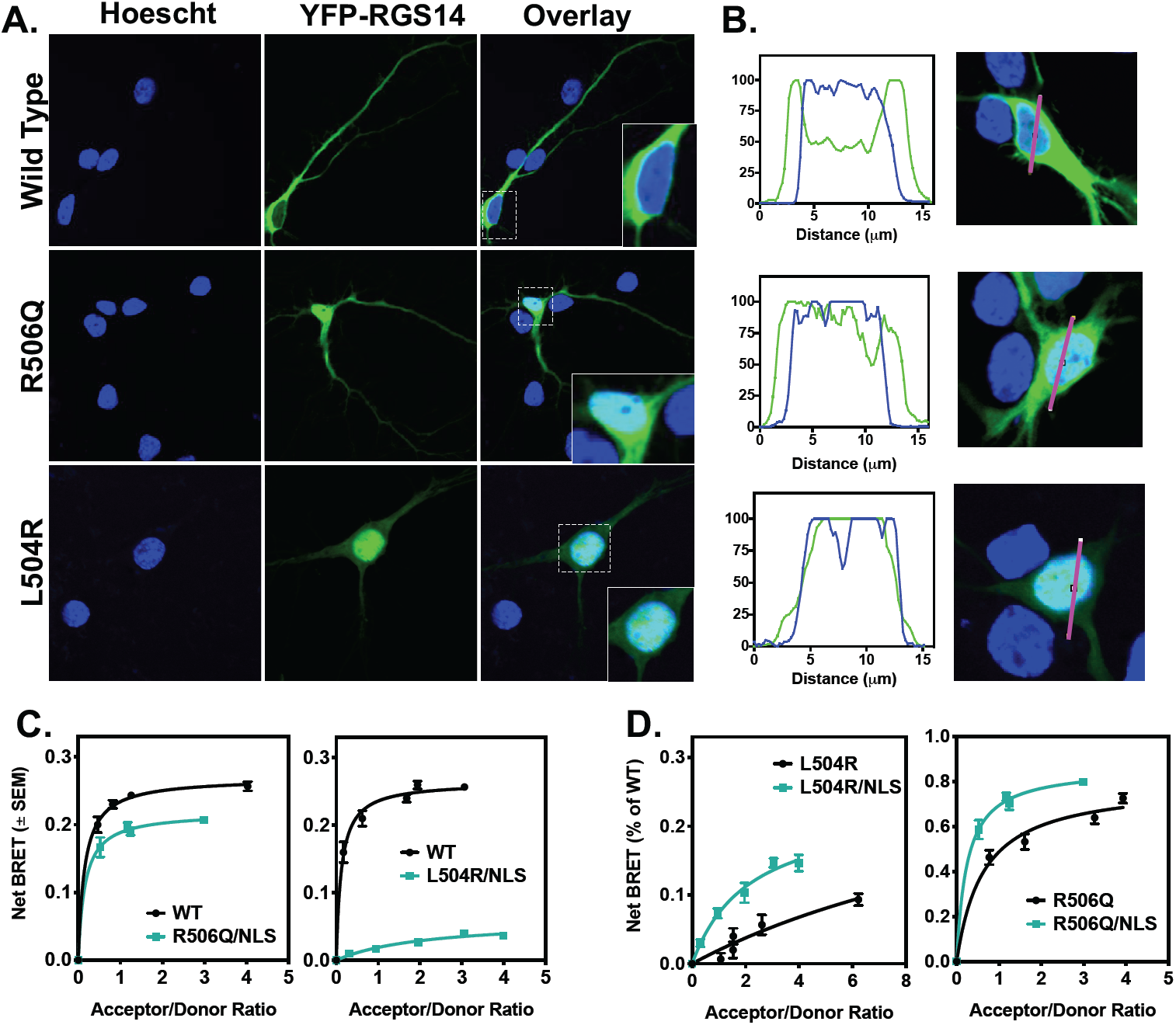
Human variants LR and RQ disrupt RGS14 nucleocytoplasmic equilibrium to favor the nucleus over the cytoplasm. Primary hippocampal neurons were infected with AAV-eYFP-RGS14 or RGS14 carrying variants LR or RQ for 18 hours and then imaged using live cell confocal microscopy. **A)** Wild type RGS14 does not co-localize with Hoechst in neurons under unstimulated conditions. The RQ variant occupies both the nucleus and the cytoplasm, while the LR variant localizes entirely to the nucleus. **B)** Quantification of YFP (RGS14) and Hoechst overlay for WT RGS14 and each of the variants provides a measure of co-localization. Approximately 10 images were collected per condition and representative images are shown. **C)** RGS14-Luc and Gαi1-eYFP-GDP interactions in live HEK cells were measured as static Net BRET. Since the RGS14 NES motif overlaps with the GPR binding site with Gαi1-GDP, an NLS mutation (NLSm) was introduced to prevent nuclear localization. Thus, any change in net BRET from wild type is due to effects of Gαi1-GDP binding, independent of mislocalization to the nucleus and interaction with XPO1. Net BRET for both RGS14 RQ (n=3) and RGS14 LR (n=3) is partially (although not fully) restored when an NLS mutation is introduced. **D)** Comparison between L504R to L504R/NLSm, and R506Q to R506Q/NLSm (L504R and R506Q data from Figure 2). Adding the NLS mutation enhances Gαi1-GDP association, indicating that XPO1 and Gαi1-GDP interactions are both impacted by the L504R and R506Q mutations.

Given that the RGS14 binding motifs for XPO1 and Gαi1-GDP overlap, coupled with previous studies showing that Gαi1-GDP interactions can affect nuclear localization (25), we next investigated whether the reduction in RGS14-Luc:Gαi1-YFP binding observed in Figure 2 was due to RGS14 sequestration in the nucleus, or if there was an additional inhibitory effect on the RGS14:Gαi1 binding interaction itself. For these studies, we generated mutations that disrupt the NLS motif (R208A/K209A/K210A; NLSm) (Figure S4) in combination with our RGS14-Luc LR and RGS14-Luc RQ constructs to make RGS14-Luc LR/NLS and RGS14-Luc RQ/NLS, respectively. These double mutant constructs are unable to translocate to the nucleus (Figure S4). Therefore, contributions of loss of XPO1 binding and nuclear sequestration should be eliminated, allowing a measure of the effect of these human variants on RGS14-Luc:Gαi1-YFP binding alone. We found that RGS14 LR/NLS and RGS14 RQ/NLS partially, but not fully, restored Gαi1-YFP binding (Figure 4C). Comparing LR to LR/NLS and RQ to RQ/NLS as a percentage of wild type (Figure 4D) using the data generated in Figure 2C shows an incomplete rescue of RGS14 binding to Gαi1-YFP in both the LR and RQ variants when an NLS mutation was added (Figure 4C), suggesting at least a partial disruption of RGS14 binding to Gαi1-GDP unrelated to XPO1 binding. Furthermore, these results suggest that any RGS14 LR or RGS14 RQ that is translocated from the nucleus will be an unlikely target of Gαi1-GDP, thus further disrupting plasma membrane recruitment and proper cellular trafficking necessary for inhibitory actions on LTP (Figure 1). Together, our BRET data (Figure 2) and imaging data (Figure 4) indicate that naturally occurring rare human variants LR and RQ confer aberrant cellular trafficking properties to RGS14.

### The XPO1 binding motif in RGS14 fits a class 3 model

RGS14 is a nucleocytoplasmic shuttling protein in neurons that utilizes XPO1 as its export receptor (Figure 3), and variants LR and RQ disrupt this interaction (Figure 4). To explore the molecular basis for this, we next created a structural model of the RGS14 NES motif bound to XPO1 (Figure 5). XPO1, also known as CRM1, recognizes a diverse range of NES motifs that can be grouped into ten classes according to their unique spacing of hydrophobic residues (39). The RGS14 NES motif embedded within the GPR motif forms an all α-helical secondary structure that matches a Class 3 spacing motif: ϕ_1_XXϕ_2_XXXϕ_3_XXϕ_4_ where ϕ is L, V, I, F, or M, and X is any amino acid (Figure 5A). We therefore hypothesized that the RGS14 NES would fit a class 3 binding model to XPO1. The RGS14 GPR peptide, which contains the full NES, was crystallized previously in complex with Gαi1-GDP (35), and a class 3 NES (mDia2) was recently crystallized with XPO1 (39). Using this information, we generated an RGS14-NES/XPO1 model by structurally aligning RGS14’s NES with the crystal structure of the mDia2 NES–XPO1 complex (Figure 5B). The alignment of the two NES-containing helices reveals optimal positioning of the hydrophobic residues (ϕ) necessary for interaction with XPO1 (Figure 5C). These ϕ residues stabilize the RGS14–XPO1 interaction through hydrophobic interactions with XPO1 pockets P0-P3, mirroring the previously reported mDia2-XPO1 complex (Figure 5D) (39). When co-expressed in HEK293 cells, co-immunoprecipitation of RGS14 with Flag-XPO1 confirmed that RGS14, but not RGS14 NESm, is a binding partner of XPO1 (Figure 5E).

**Figure 5.**
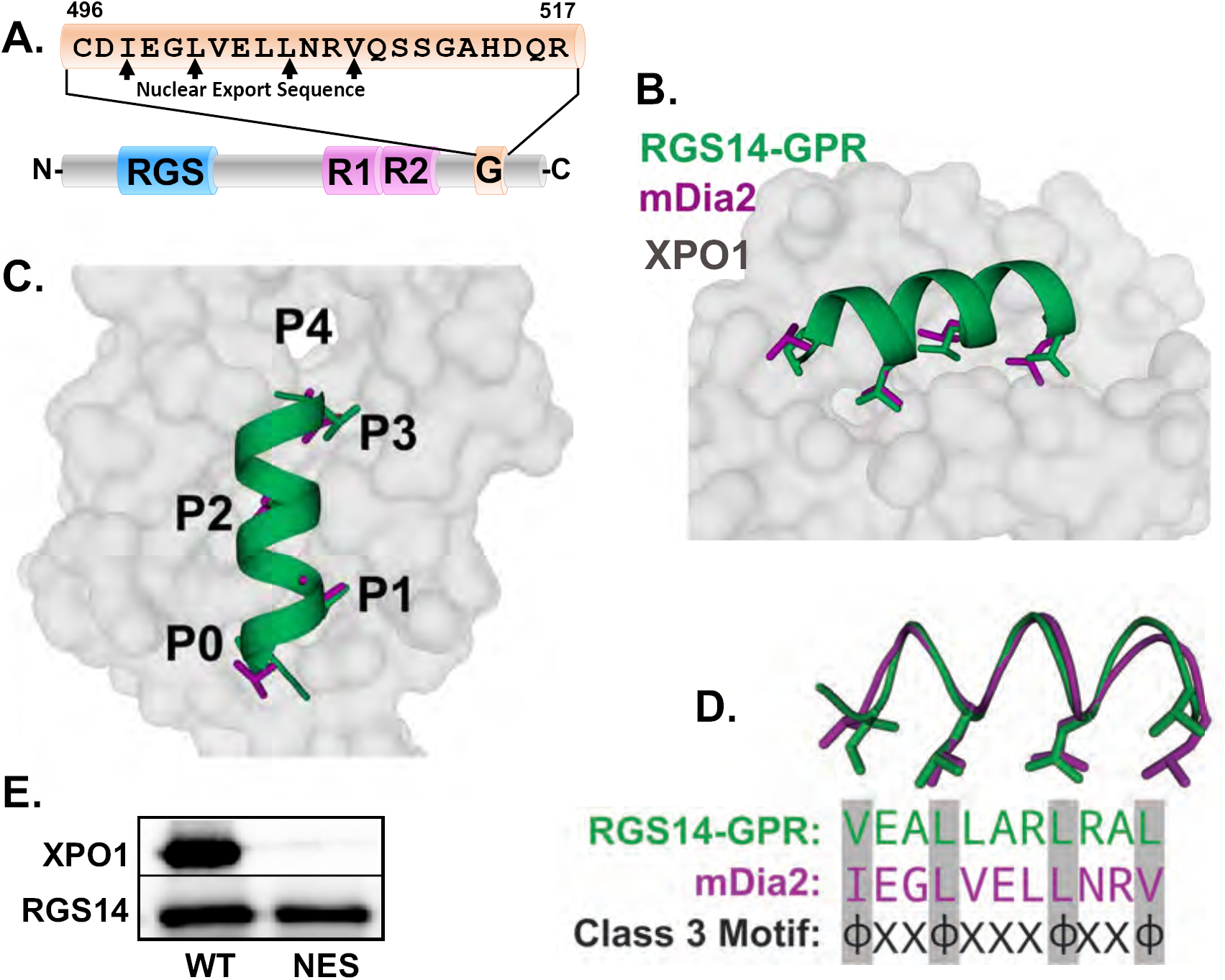
The GPR motif of RGS14 contains an embedded class 3 nuclear export sequence that binds XPO1. Structural modeling of the RGS14 GPR/NES motif binding to Exportin 1 (XPO1). **A)** The NES of RGS14 is embedded within the GPR motif and follows a class 3 pattern: ϕ_1_XXϕ_2_XXXϕ_3_XXϕ_4_. **B)** The crystallized GPR peptide of RGS14 (PDB 1KJY) was structurally aligned with a similar class 3 NES motif, (mDia2 PDB 5UWP) crystallized with XPO1. **C-D)** The RGS14 NES fits a class 3 binding motif, occupying pockets 0-3, but not pocket 4, as reported previously (39). **E)** RGS14 WT, but not RGS14 NES, co-immunoprecipitates with Flag-XPO1. Flag-XPO1 and RGS14 WT or NES were transfected into HEK cells and Flag was immunoprecipitated (Flag affinity gel). RGS14 (1:1000) and XPO1 (1:300) protein were measured by immunoblot.

### Human variants LR and RQ disrupt RGS14 interactions with XPO1

As shown above, RGS14 is a nuclear shuttling protein that utilizes XPO1 as a nuclear export receptor, and human variants RQ and LR cause RGS14 to mislocalize and accumulate in the nucleus. Based on this, we hypothesized that these variants disrupt RGS14 binding with the XPO1. To test this idea, we examined whether these variants altered RGS14 complex formation with purified XPO1 and its obligate binding partner, Ran-GTP. Purified bacterially expressed recombinant RGS14 WT, RGS14 NES, RGS14 LR or RGS14 RQ were mixed with purified XPO1-GST and constitutively active His-Ran Q69L-GTP. After mixing for 2 hrs for complex formation to occur, His-Ran QL-GTP was recovered by Ni-NTA affinity pull-down along with XPO1-GST and/or RGS14 in complex. As expected, RGS14 WT was precipitated in complex with XPO1. In contrast, RGS14 NES, RGS14 LR, or RGS14 RQ did not bind XPO1 (Figure 6A-B). We next utilized the RGS14-NES/XPO1 model described above (Figure 5) to further explore the mechanism by which these variants disrupt RGS14 binding to XPO1. As the reported RGS14 GPR crystal structure utilized rat RGS14 (35), the human variants R507Q and L505R correspond to rat residue number R506Q and L504R. Based on our model, arginine 506 (R506) in wild-type RGS14 contains a positively charged nitrogen atom that is 5.5 Å away from a negatively charged oxygen atom of glutamic acid in XPO1, which is in line with a possible ionic interaction. We then mutated this residue in our model to the RQ variant, glutamine 506 (Q506), in order to provide mechanistic insight for the decreased nuclear export present with the RQ variant (Figure 6C). The variant (Q506) disrupts the ionic interaction that was present between R506 and glutamic acid in XPO1. The distance between Q506 and the glutamic acid in XPO1 is too great for hydrogen bonding, suggesting this variant will reduce affinity of RGS14 for XPO1 (Figure 6C) and thus explains the mixed phenotype. Although RGS14 RQ can bind XPO1, it does so less efficiently than WT RGS14. We next wanted to visualize the impact of the LR variant, arginine 504 (R504). In wild-type RGS14, leucine 504 (L504) fits nicely within the P2 hydrophobic pocket of XPO1, stabilizing the complex through hydrophobic interactions. The LR variant (R504) places a charged arginine within the P2 hydrophobic pocket of XPO1 causing a polar incompatibility that will greatly disrupt the RGS14-XPO1 interaction (Figure 6D-E). This severe mutation within the binding interface of RGS14 and XPO1 suggests the LR variant is highly unlikely to interact with XPO1 and thus explains why RGS14 LR becomes trapped within the nucleus.

**Figure 6.**
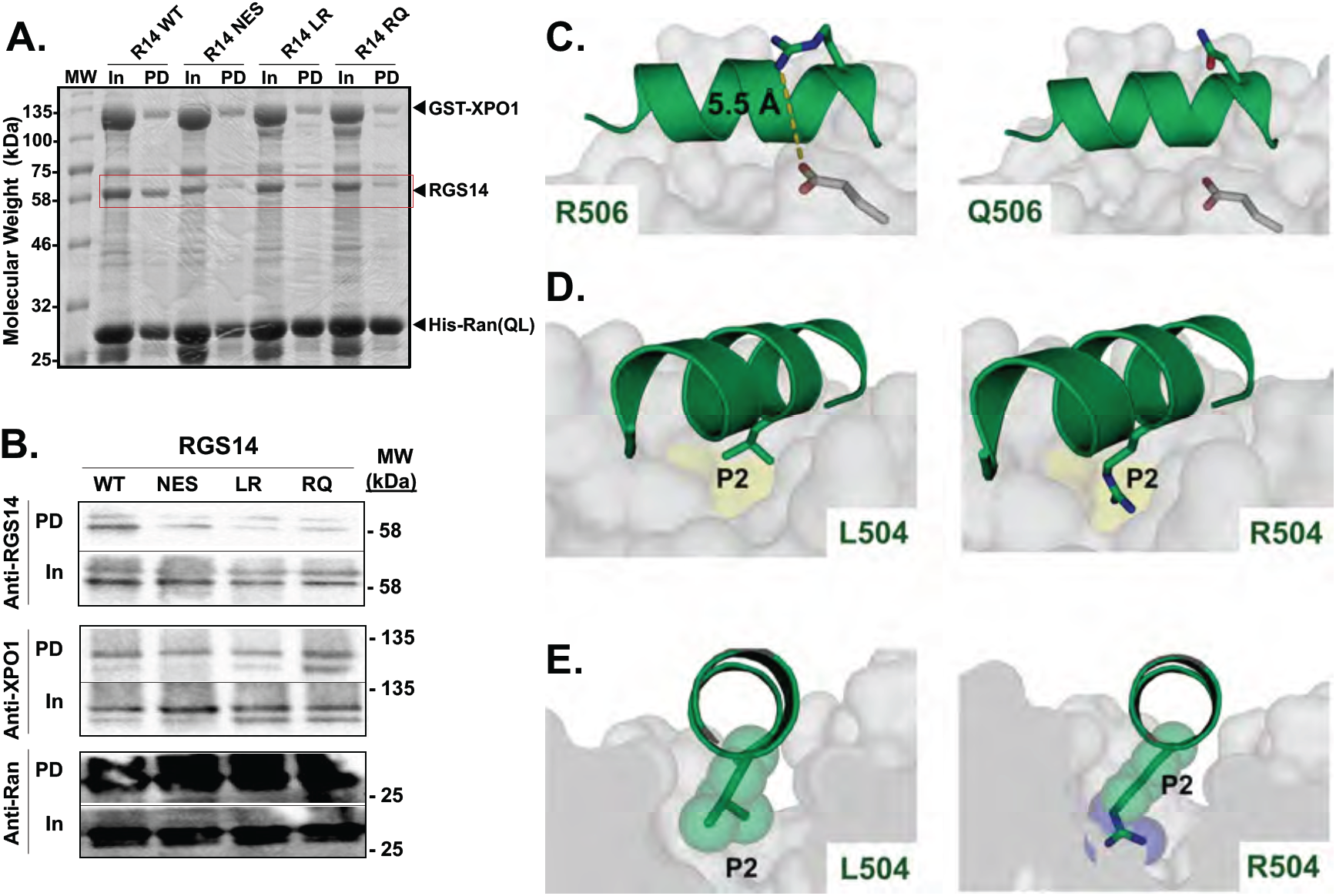
Human variants LR and RQ disrupt RGS14 binding to Exportin 1 (XPO1) through loss of side chain interactions. **A-B)** Recombinant RGS14 WT and variants LR and RQ were purified from E. coli and mixed with purified GST-XPO1 and constitutively active His-Ran (GV)-GTP. His-Ran (G/V)-GTP was captured by Ni-NTA affinity pull-down and complexes were assessed by Coomassie **(A)** and immunoblot **(B)**. WT RGS14 was recovered in complex with XPO1, but RGS14 NES, RGS14 LR, and RGS14 RQ did not bind XPO1. Blots are representative of 5 independent experiments. **C)** Structural modeling of the R506Q variant binding to XPO1. Left panel is the wild type amino acid, arginine, which makes a salt bridge with a nearby glutamate near the binding pocket on XPO1. Mutation to a glutamine (right panel) removes this salt bridge, presumably destabilizing the protein-protein interaction. **D-E)** Structural modeling of the L504R variant onto XPO1. The leucine interacts directly with the hydrophobic P2 pocket (left panel). Mutation of this amino acid to an arginine (right panel) introduces a charged side chain into a hydrophobic pocket. These data mechanistically explain the difference in phenotype (R506Q being a subtler phenotype compared to L504R).

### Human variants LR and RQ disrupt RGS14 suppression of LTP in hippocampal slices

The profound disruption of RGS14 binding to Gαi1-GDP and XPO1 observed with the LR and RQ human variants, which results in their subcellular mislocalization, predicts that these variants will negatively impact RGS14 suppression of synaptic plasticity. To examine this, we utilized the experimental system of RGS14 ectopic expression in CA1 hippocampal slices as shown in Figure 1 and as described previously (21). We generated wild type (WT) AAV-YFP-RGS14 and AAV-YFP as positive and negative controls and compared these to AAV-YFP-RGS14 LR and AAV-YFP-RGS14 RQ. Each of these AAV constructs were exogenously expressed in area CA1 of cultured hippocampal slices (Figure 7A). We found that wild type WT YFP-RGS14 was detected throughout the soma, dendrites, and spines (Figure 7A), consistent with expression patterns in dissociated hippocampal neurons (Figure 4A). YFP-RGS14 RQ filled the entire cell, occupying both the nucleus and cytoplasm, while YFP-RGS14 LR localized to the nucleus (Figure 7A), consistent with our findings in dissociated neurons (Figure 4A). We recorded excitatory post synaptic currents (EPSCs) at baseline and following an LTP induction protocol, as before (Figure 1 and (20)). Neurons from slices with YFP expression alone showed robust LTP (Figure 7B), while WT YFP-RGS14 expression completely blocked LTP, as expected (Figure 7C). Consistent with our other data here, the YFP-RGS14 RQ variant allowed induction of a short lasting potentiation (Figure 7D), while the YFP-RGS14 LR variant blocked RGS14’s capacity to inhibit LTP (Figure 7E). Together, these findings demonstrate that: 1) RGS14 mislocalization is associated with impaired postsynaptic plasticity functions; and 2) the degree of nuclear accumulation is correlated with the degree of RGS14 dysfunction as a regulator of postsynaptic plasticity.

**Figure 7.**
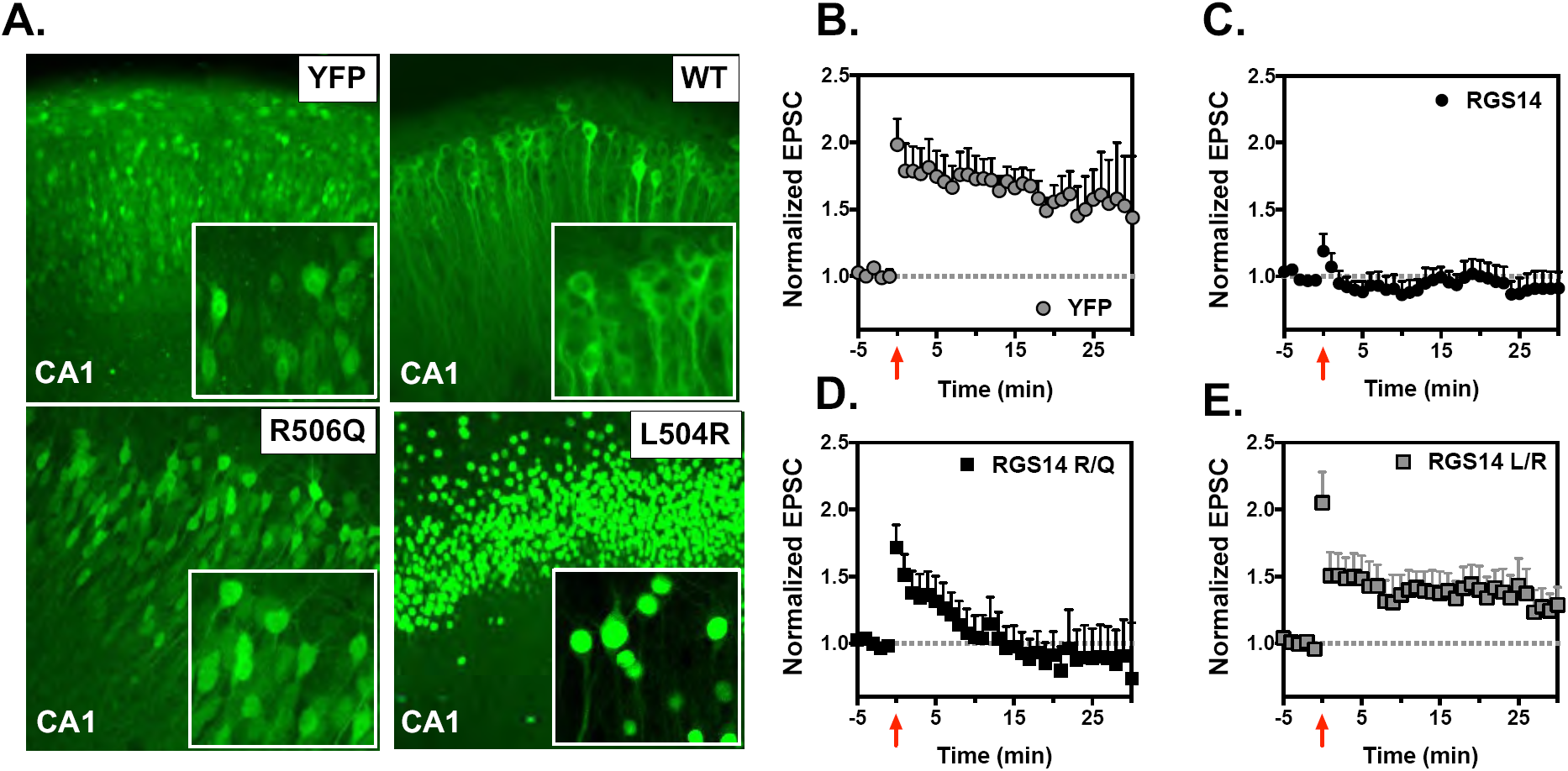
Human variants LR and RQ that disrupt RGS14 nuclear equilibrium also block RGS14 capacity to inhibit long term potentiation in hippocampal slices. AAV-eYFP-RGS14 was used to make AAV2/9 viruses, including wild type RGS14 as well as human variants RQ and LR. Furthermore, a stop codon was generated as follows AAV-YFP-STOP-RGS14, to make a truncated YFP virus lacking RGS14. Virus was injected into mouse CA1 slice cultures and incubated for one week, at which point electrophysiological recordings were made to assess LTP. **A)** Expression of YFP-RGS14 in CA1 brain slices. WT RGS14 fills the cytoplasm of neurons. RQ fills both the cytoplasm and nucleus, while LR localizes predominantly to the nucleus, entirely consistent with our data in dissociated neurons. Images are representative of 6-9 experiments, **B)** Long term potentiation (LTP) was induced by an LTP pairing protocol in CA1. YFP alone (n=8) had no effect on LTP, but in contrast, WT RGS14 (n=6) completely inhibits LTP in CA1, a hippocampal region where RGS14 is not natively found. Potentiation lasting only about 10 mins was induced in RQ variant-expressing neurons (n=9), whereas the LR variant (n=9) failed to suppress LTP. This suggests that the degree of mislocalization by these variants correlates with ablation of RGS14 function in CA1 neurons. Data is presented as mean +/-SD.

### Variant LR disassociates RGS14 actions on synaptic plasticity from spatial learning in mice

We previously reported that genetic loss of RGS14 (RGS14 KO mice) restores LTP in hippocampal area CA2 that is correlated with enhanced spatial learning as measured by the Morris Water Maze test (20). Based on our observation that the LR variant completely ablated RGS14-dependent suppression of LTP (Figure 7), we hypothesized that RGS14 LR may behave as a functional knock out and recapitulate the enhanced spatial learning phenotype of the RGS14 KO mice (20). To test this idea, we generated a novel mouse line carrying the RGS14 LR variant using CRISPR/Cas9 on a C57Bl/6J background. To confirm that RGS14 LR variant mice exhibited the characteristic aberrant localization of RGS14 in hippocampal CA2 neurons, we performed immunohistochemical staining of hippocampal slices. We observed that, like our findings with cultured hippocampal slices (Figure 7) and neurons (Figure 4), homozygously expressed RGS14 LR was localized to the nuclei of CA2 neurons as compared to RGS14 WT, which exhibited typical diffuse staining throughout the soma and neurites of CA2 neurons but was absent from nuclei (Figure 8A). Heterozygously expressed RGS14 LR showed a mixed expression pattern present in soma, neurites and the nuclei of CA2 neurons. These findings with whole animal are completely consistent with our observations in cultured hippocampal slices (Figure 7) and primary neurons (Figure 4).

**Figure 8.**
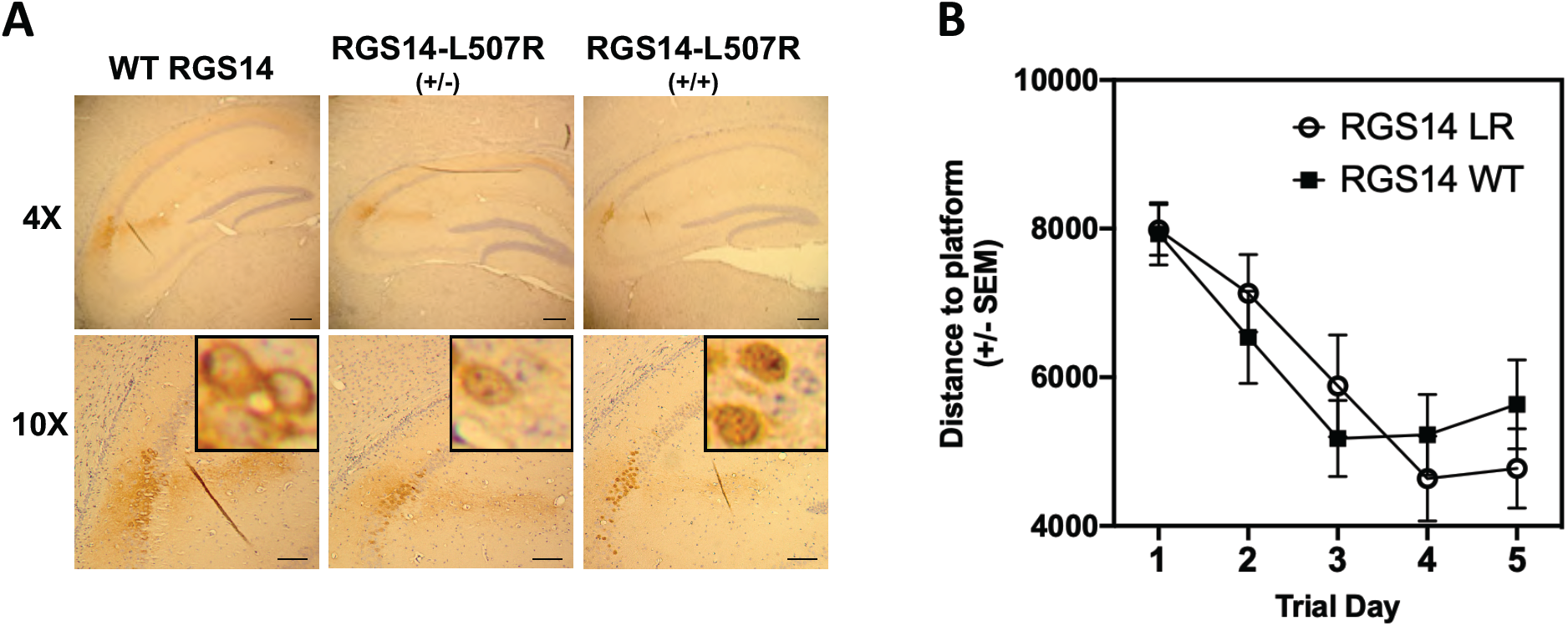
The RGS14 variant LR localizes to the nuclei of hippocampal neurons but does not alter hippocampal-based spatial learning in CRISPR/Cas9 mice. Mice carrying the LR variant were generated by CRISPR/Cas9 (RGS14 LR mice). **A)** Expression and localization of RGS14 LR in CA2 hippocampal neurons was confirmed by immunohistochemical staining. Consistent with our slice electrophysiology data, RGS14-LR was predominantly nuclear, while RGS14 WT was localized to soma, dendrites, and spines but excluded from the nucleus of CA2 hippocampal neurons. The heterozygous mice exhibited a mixed nuclear-cytoplasmic phenotype. **B)** RGS14-LR mice (n=20) and RGS14 WT (n=20) littermates were subjected to the Morris water Maze test. RGS14-LR mice do not exhibit enhanced spatial learning, suggesting a unique nuclear role of RGS14 on suppression of spatial learning.

We took advantage of these animals to examine RGS14 protein expression in brain regions outside of the hippocampus, particularly brain regions recently reported in monkey and human to express RGS14 (28,31). Consistent with RGS14 protein expression patterns in primates, we observed RGS14 protein expression in the central amygdala, striatum, and piriform cortex of the RGS14 LR mouse (Figure S5). RGS14 expression had not been reported previously in striatum and amygdala of rodents, due to low mRNA expression (Allen Brain Atlas: https://portal.brain-map.org/). In the RGS14 LR animals, RGS14 is concentrated in the nuclei resulting in a bright punctate staining pattern, greatly improving detection and visualization of RGS14. The concentration of RGS14 LR in the nucleus allows for effortless distinction between cells that express RGS14 and cells that do not, but which are contacted by RGS14-containing neurites.

As reported previously, RGS14 KO mice exhibit an enhancement of spatial learning as measured by the Morris Water Maze test (20). To determine the effects of nuclear-bound RGS14 LR on spatial learning, we subjected RGS14 WT and RGS14 LR mice to the Morris Water Maze test (Figure 8B). We expected the LR mice to mimic the RGS14 KO mice and exhibit enhanced spatial learning as before (20). However, quite unexpectedly, RGS14 LR mice did not exhibit an enhancement of the spatial learning phenotype and behaved similarly to WT littermates (Figure 8B). In summary, RGS14 LR mimicked one aspect of genetically ablated RGS14 (RGS14 KO), i.e. a release on the block of LTP in hippocampal neurons, but failed to recapitulate the other phenotype of RGS14 KO mice, i.e. enhancement of spatial learning (20). Taken together, these findings suggest that nuclear RGS14 may serve a distinct cellular role from RGS14 localized to postsynaptic dendritic spines.

Although RGS14 is a well-established nucleocytoplasmic shuttling protein, nuclear roles for RGS14 remain unknown. In an initial attempt to define a nuclear role for RGS14, we examined if RGS14 nuclear shuttling and localization impacted mRNA expression levels or patterns in hippocampal neurons using RNA-seq methodology (40). Cultured primary hippocampal neurons were infected with AAV expressing RGS14 with normal nucleocytoplasmic shuttling (WT), RGS14 excluded from the nucleus (NLSm), or RGS14 targeted exclusively to the nucleus (NESm) (Figure S4). After 2 weeks, neurons were harvested, mRNA was isolated and converted to sequencing library for analysis. No differences in the gene expression patterns or total number of genes expressed (10,324 of 17,324 transcripts) were observed in hippocampal neurons expressing either RGS14 exclusively in the cytosol (NLSm), the nucleus (NESm), or both (WT) (Figure S6).

## Discussion

Here, we provide evidence for the interdependence of RGS14 subcellular localization and protein function in host neurons, specifically driven by genetic diversity. Our findings suggest distinct roles for RGS14 in the nucleus and dendritic spines of hippocampal neurons, and highlight the underappreciated fact that rare human genetic variants can alter the function of affected proteins to markedly impact cellular physiology in unexpected ways.

We identified the GPR motif as being critical for RGS14 functions in hippocampal neurons. Numerous reports have shown that RGS14 is a dynamically regulated protein which exhibits a nuclear-cytoplasm-membrane shuttling phenotype favoring cytoplasmic localization in resting cells (25-27,31,36). The C-terminus of RGS14 harbors the GPR motif with an embedded NES that governs the intracellular dynamics of RGS14, and dictates its availability to interact with signaling partners necessary to carry out its many functions (41-44). A well characterized RGS14 function is regulation of synaptic plasticity in area CA2 of the hippocampus. Long term potentiation (LTP) and synaptic plasticity are mediated by a complex network of signaling events that drive global changes in connectivity, learning, and behavior. For this network to function properly, signaling proteins must be in the right place, at the right time, and for the right amount of time. RGS14 is a natural suppressor of LTP and postsynaptic plasticity in hippocampal area CA2 (20,21) that relies on dendritic spine localization to perform these functions. Here we demonstrate that genetic variants within the GPR/NES motif disrupt RGS14 localization to dendritic spines and capacity to inhibit LTP, serving to highlight the relationship between spatial localization and function for RGS14.

One surprising result from our studies is that the nuclear-bound RGS14 LR variant presented a discordant behavioral phenotype compared to the RGS14 KO phenotype in mice. We expected the RGS14 LR mice to behave identically to the RGS14 KO mice; however, that was not the case. Previous work shows that the RGS14 KO mice exhibit unrestricted LTP in hippocampal slices that is correlated with an enhancement in spatial learning (20). Whereas the RGS14 LR mice exhibited a release of the brake on LTP phenotype in hippocampal slices, they did not exhibit altered spatial learning. Reasons for this distinct phenotype are unclear. Of note, early LTP lasts 1-2 hours and is dependent on local signaling and local protein synthesis at the spines and dendrites (45,46). Both the KO and LR mice lack RGS14 in dendritic spines, resulting in unrestricted early LTP. By contrast, late LTP and linked memory last much longer and are reliant on synthesis of *new* mRNA/protein from the nucleus (47,48). Hippocampal-based spatial learning is dependent on consolidation of these late LTP processes, including nuclear-synapse shuttling (48,49). In the RGS14 KO mice, RGS14 is neither at the spines nor in the nucleus, whereas RGS14 LR exists primarily in the nucleus. In the case of RGS14 LR mice, any enhancement of early LTP at the spines may not be properly consolidated in late (nuclear-dependent) LTP processes due to undefined actions of RGS14 in the nucleus. Conversely, RGS14 KO mice have a release of molecular brakes at the spines and dendrites, *and RGS14 is absent from the nucleus*, allowing for unabated inhibition of plasticity thereby suggesting a possible distinct nuclear role of RGS14.

Specific roles for RGS14 in the nucleus remain a mystery. Previous studies with HeLa cells reported that recombinant RGS14 translocates from the cytosol to the nucleus following mild heat stress (26). There, RGS14 localized to promyelocytic leukemia (PML) bodies, which have been implicated in cellular processes such as apoptosis, transcriptional regulation, DNA repair and replication (50). However, we have not observed native RGS14 localization at PML bodies in mouse hippocampal neurons, monkey striatal neurons (31), or rat B35 neuroblastoma cells (27), suggesting RGS14 PML body localization may be limited to unnatural host or non-neuronal cells. RGS14 lacks a DNA-binding motif but may nonetheless regulate the expression of synaptic proteins critical for late term LTP and memory in neurons. This could occur through RGS14 interactions with accessory proteins, chromatin remodeling, mRNA splicing machinery, or other nuclear processes. Consistent with this idea, native RGS14 has been observed to exist at intranuclear membrane channels, and within both chromatin-poor and chromatin-rich regions of the nucleus of a neuronal cell line (27). In addition, a subset of nuclear RGS14 localizes adjacent to active RNA polymerase II. Together, these findings suggest potential roles for RGS14 in transcriptional regulation (27,51,52).

Even so, our pilot studies here with cultured hippocampal neurons showed no differences in the number or variety of mRNA transcripts in neurons expressing either normal RGS14, RGS14 excluded from the nucleus, or RGS14 trapped in the nucleus (Figure S6). Of note, the neurons in this study were at rest and in culture. By contrast, proper RGS14 nuclear targeting and actions may require specific postsynaptic activity in intact hippocampal slices or animals. Though speculative, a possibility is that synaptic activation could redirect a subpopulation of RGS14 and/or other regulatory factors to the nucleus to modulate transcription. RGS14 is not the only RGS protein found in the nucleus (53,54). RGS14’s closest relatives, RGS10 and RGS12, as well as certain other RGS proteins have been reported to localize to the nucleus (55-57), with limited understanding of their functions there. It is quite possible that these RGS and RGS14 only impact gene expression in response to specific stimuli. As we show here, human variants target RGS14 to the nucleus constitutively, which would disrupt any temporal regulation or necessary post-translational modification for RGS14 necessary for nuclear functions and would confer a different phenotype than would be exhibited by normal nuclear shuttling of RGS14.

Another possible nuclear role for RGS14 may involve synapse-nucleus signaling, which is important for spatial memory (58) and has been postulated to facilitate synapse-specific targeting of newly synthesized mRNAs or proteins following an LTP stimulus (59-62). Signals from the activated synapse must reach the nucleus, and newly synthesized mRNA must reach the activated synapse within a relatively short window in order to support synaptic strengthening and maturation (60). How the neuron coordinates this has become generally accepted under the Synaptic Tagging and Capture (STC) hypothesis (60,63). RGS14 could play a role in either retrograde or anterograde trafficking, coordinating signals from the spine to the nucleus or coordinating cargo delivery back to the spine. In support of this idea, RGS14 has been found to colocalize with cytoskeletal elements, including microtubules, and influences tubulin polymerization (27,64,65), which enables trafficking of cargo to the tagged synapse (66,67). As a multi-functional scaffold, RGS14 integrates G protein, Ca^++^/CaM/CaMKII, H-Ras/ERK, and 14-3-3γ signaling (17,18,21,36,68,69), all critical mediators of both early and late stage LTP and learning (70-73), and thus could be acting as a scaffold to coordinate signaling between synapse and nucleus. Importantly, while some studies suggest that proteins at the activated synapse translocate to the soma/nucleus, others indicate that action potentials stimulate nuclear import of somatic molecules to mediate transcription (59), suggesting a nuclear/somatic pool of RGS14 could have distinct functions from a spino-dendritic pool of RGS14.

The simplest explanation for RGS14 targeting to the nucleus is to redirect and sequester the protein away from the spine as a mechanism by the neuron to remove the brake on LTP, with no specific nuclear function (i.e. time-out). Upon LTP induction, the spine undergoes massive remodeling, including substrate phosphorylation, AMPA receptor trafficking, and actin cytoskeleton rearrangement (74). As mentioned above, RGS14 engages multiple binding partners strongly linked with regulation of LTP at the spine (12-15,17,69,74,75). Thus, RGS14 pre-positioning at the spine allows for immediate participation in, and regulation of, these processes. Removing the available pool of RGS14 from spines may be one way in which the cell can allow for differential regulation of these LTP-linked pathways. Although our preliminary RNA-seq findings did not identify target genes modulated by RGS14, the discordant phenotype of nuclear-bound RGS14 LR versus RGS14 KO suggests that RGS14 is not simply passively hiding out in the nucleus, but likely confers an expression- and localization-dependent outcome on nuclear processes necessary for spatial learning. Ongoing and future studies will explore these possibilities.

We recently reported the native expression of RGS14 in human and monkey brain, demonstrating the presence of RGS14 protein in unexpected brain regions and in the nuclei of certain neuronal populations (31). In addition to the hippocampus, RGS14 is robustly expressed in the piriform cortex, caudate, putamen, globus pallidus, substantia nigra, and amygdala of primate brain. Our studies here confirm for the first time in adult rodents the expression of RGS14 in brain regions outside of the hippocampus, including striatum and amygdala, and raise questions about RGS14 roles beyond the hippocampus. One feature these brain regions all share is experience-based synaptic plasticity (76-78), suggesting that RGS14 may serve as a broad regulator of plasticity in these brain regions. Within the striatum, RGS14 could be regulating changes in learned behavior linked to reward and addiction (79). Other RGS proteins have been implicated in various measures of drug seeking and drug reward behavior (80). For example, both RGS9-2 and RGS7 in the striatum modulate locomotor responses to cocaine (81,82). Within the amygdala, RGS14 could be regulating fear memory and stress responses (78). Consistent with this idea, a recent report showed that a global loss of RGS14 (RGS14 KO) enhances certain fear behaviors in female mice (83). Although the report demonstrates this behavioral phenotype is linked to the known RGS14-rich CA2 region of the hippocampus, this study cannot rule out contributions of RGS14 within the central amygdala, a region long known to mediate behaviors associated with fear. While we do not yet know the significance of RGS14 in these regions, these questions motivate current and ongoing studies.

Our findings here were made possible by the study of natural genetic variation within the human RGS14 gene. We examined 13 human variants within the GPR motif and found two that markedly altered RGS14 function in unexpected ways distinct from the “normal” human reference sequence. Genes/proteins are typically studied as canonical reference sequences (defined as normal) that do not reflect the rich genetic diversity in the human population. Our findings here show that this genetic diversity can have profound impacts on protein function and linked physiology. The variants chosen for study were selected from a dataset of human variants within the RGS protein family (4). Here we describe a process for characterizing such variants, whereby a functional domain is isolated and then screened for dysfunction when variants are introduced. We find that, while these variants were derived from a “healthy” population, they may impart important functional consequences in human carriers that would not be detected by studying a monogenic disease model. As an example, genetic variants within the non-coding region of RGS16 are linked with self-reported “morning people” (84,85), an intriguing human trait that would not otherwise be captured by studies focused exclusively on disease-linked SNPs.

Regulatory proteins, such as those in the RGS family, that contribute to complex signaling cascades (15,86,87) are more likely to contribute to complex polygenic diseases or physiological traits, particularly in combination with variants in other proteins. As shown here, RGS14 protein harboring rare variants LR or RQ could lead to aberrant signaling in synaptic plasticity in striatum or amygdala, for example contributing to a predisposition to fear learning or addiction, among others. Within the hippocampus, where RGS14 function is best characterized, variants LR and RQ tip the nucleo-cytosolic-membrane shuttling balance in favor of the nucleus, leading to functional consequences in RGS14-linked plasticity. Indeed, protein mislocalization due to a coding variant has been shown to contribute to protein dysfunction in other complex diseases (88), underscoring the importance of signaling proteins being in the right place, at the right time, for the right amount of time for proper function.

In summary, we show that RGS14 harboring human genetic variants exhibits aberrant subcellular trafficking and mislocalization that results in loss of function in dendritic spines. Based on these observations, we propose a working model suggesting that RGS14 regulates multiple processes for both early and (potentially) late LTP at the spines and nucleus, respectively (Figure 9). At postsynaptic spines, RGS14 regulates GPCR/G protein, H-Ras/ERK, Ca^++^/CaM and other signaling events to suppress LTP (15,18,19,75) (Figure 9A). Mislocalization and nuclear accumulation of RGS14 due to rare genetic variant LR (or variant RQ, to a lesser extent) leads to absence of RGS14 from spines and dendrites, rendering the protein unable to modulate early LTP signaling (Figure 9B). Overall, our findings here highlight the importance of dynamic spatial regulation of signaling proteins in the context of synaptic plasticity, and demonstrate the contribution of naturally occurring rare genetic variation to RGS14 compartmental equilibrium. These data underscore the importance of considering not only the function of proteins derived from “canonical” sequences, but also those representing diverse human populations. The observation of RGS14 expression in “plastic” regions outside of the hippocampus, such as the amygdala and striatum, raises questions about RGS14 control of plasticity within and behaviors associated with these brain regions. Whether these genetic variants confer the expression of unique traits in human carriers remains an engaging question meriting further study within the context of hippocampal-based memory and beyond.

**Figure 9.**
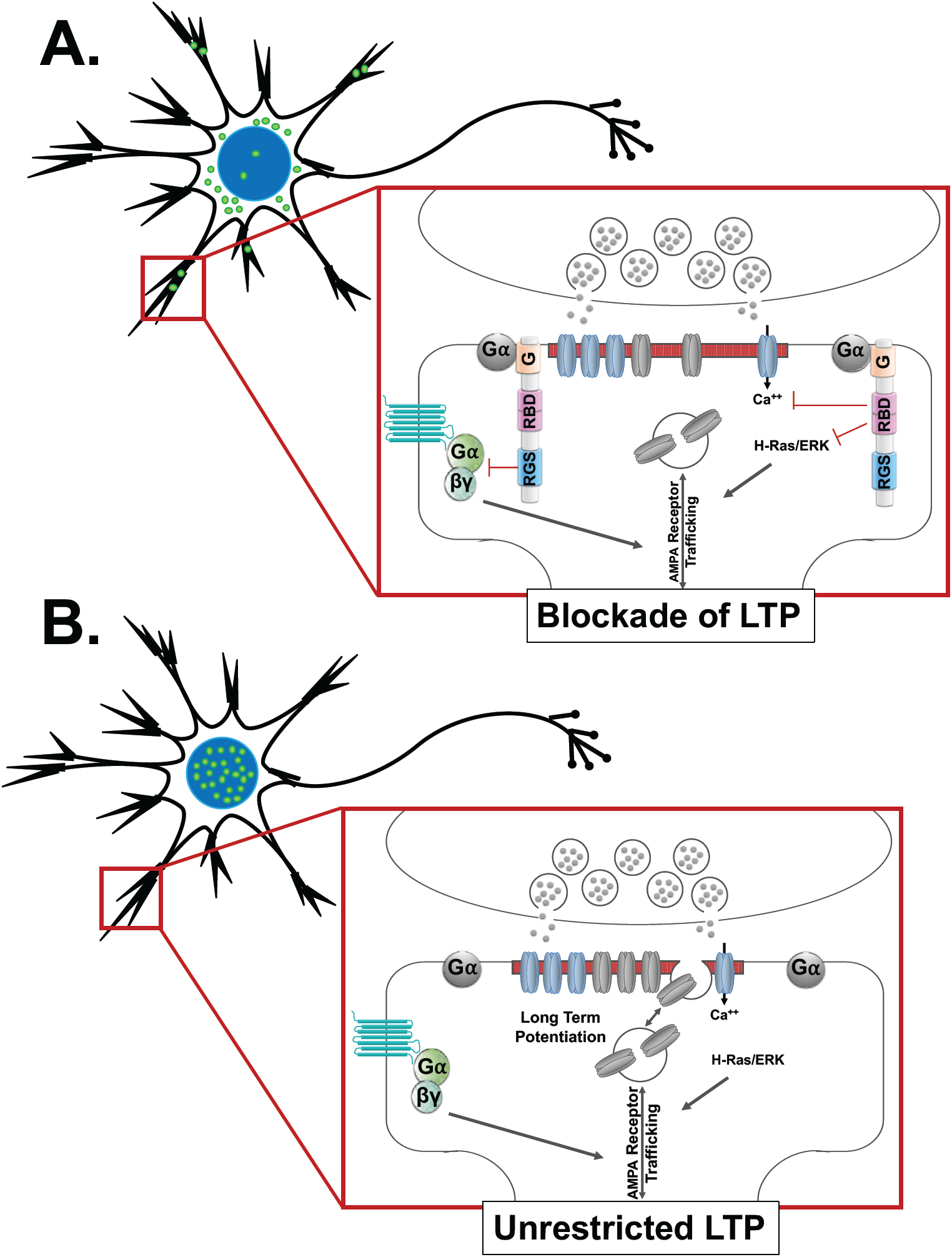
Proposed working model of RGS14 spatial-dependent inhibition of LTP. **A)** Wild type RGS14 (green dot) is disperse throughout the neuron, occupying multiple compartments including the nucleus, the cytoplasm, and the dendrites and spines (red box). Within a dendritic spine (expanded red box), Gα-GDP interaction with the G protein regulatory motif (G) brings RGS14 to the micro-compartment necessary for inhibition of GPCR signaling by the RGS domain, inhibition of H-Ras/ERK signaling by the Ras binding domains (RBD), and inhibition of Ca^++^ signaling, all of which of which support LTP in the absence of RGS14. **B)** Variant forms of RGS14 (green dots) are sequestered in the nucleus and cannot reach dendrites (red box). Due to their mislocalization, they cannot interact with, and be recruited by, Gα-GDP to dendritic compartments (expanded red box). The lack of RGS14 at these micro-compartments allows for unrestricted GPCR, H-Ras/ERK and Ca^++^ signaling, removing the block on LTP.

## Experimental procedures

### Constructs and reagents

Leptomycin B was obtained from Sigma and cells were treated at 20 nM for 2 hours. Rat RGS14 was cloned into phRLucN2 as described previously (17), pLic-GST as described previously (31), and pAAV-hSyn-YFP using restriction sites AgeI and HinDIII. pET-His-hRan-Q69L was acquired from AddGene (catalog #42048) and pGEX-TEV-hCRM1 (XPO1) was kindly provided by Dr. YuhMin Chook (UT Southwestern) (89). Flag-hCRM1 (XPO1) was obtained from AddGene (catalog # 17647). pET-Ran(Q69L) was obtained for Add Gene (catalog #42048). Isopropyl β-D-1-thiogalactopyranoside (IPTG) was purchased from Fisher Scientific. Primers used to make mutant constructs are included in Table S1. Anti-RGS14 was obtained from Proteintech. Anti-XPO1 was purchased from Sigma.

### Purification of recombinant protein and measurement of pure protein-protein interactions

RGS14 was purified as described previously (24). Briefly, MBP-TEV-RGS14 was expressed in BL21 (DE3) *Escherichia coli* (E. Coli), pellets were resuspended in 50 mM HEPES, 200mM NaCl, 10% glycerol, and 2mM BME. Bacterial lysate was passed over a glutathione affinity column, washed with PBS, eluted with 10mM reduced glutathione in 50mM Tris (pH 8), and cleaved overnight with TEV protease. Purity was verified by coomassie. His-Ran Q69L (Ran QL) was purified as follows. BL21 bacterial cells were transformed with His-Ran QL and protein expression was induced with IPTG at 37°C for 2 hours. Bacterial pellets were resuspended in 50mM HEPES, 150mM NaCl, 10% glycerol, 10mM GTP, 2mM BME, 20mM imidazole, 2mM MgCl_2_, and PMSF. Lysates were then passed over a Ni^2+^ affinity column, washed with resuspension buffer and finally eluted with resuspension buffer containing 200mM imidazole. Purity was verified by coomassie. GST-CRM1 (Exportin 1, or XPO1) was purified as described by the Chook Lab (89) as follows. BL21 bacterial cells expressing GST-CRM1 were induced with IPTG overnight at 25°C. Lysate (40 mM HEPES pH 7.5, 2 mM MgSO_4_, 200 mM NaCl, 5 mM dithiothreitol (DTT), 10% glycerol, and protease inhibitors) was then passed over a glutathione affinity column (Sepharose 4B, GE Healthcare), washed with buffer (40 mM HEPES, 2 mM MgSO_4_, 150 mM NaCl, 2 mM DTT) and beads were resuspended in PBS. CRM1/XPO1 was cleaved from the beads by incubating with TEV protease overnight at 4°C. Purity was verified by coomassie. Immunoprecipitation with FLAG and immunoblotting was performed as described previously (25,36).

### Structural modeling

The model depicting RGS14-NES/XPO1 interactions was generated in PyMol (The PyMOL Molecular Graphics System, Version 2.0 Schrödinger, LLC.) by structurally aligning the RGS14 GPR peptide/Gαi1 structure (PDB code 1KJY) with the XPO1/mDia2 structure (PDB code 5UWP). Figures were constructed using PyMol.

### Human genetic variants and lollipop plot

Human variant information was obtained from the Genome Aggregation Database (GnomAD version 2.0) at the Broad Institute (http://gnomad.broadinstitute.org) (32). Data was sorted by missense and synonymous annotation. The lollipop plot was then generated by open source code available on GitHub (90). Human variants were generated in rat RGS14 (of which the GPR motif shares 100% identity compared to human but the amino acid number is less one) and are reported in this manuscript as such. Primers used to make the human variants as well as functional mutations are listed in Table S1.

### Bioluminescence Resonance Energy Transfer (BRET)

BRET was performed as described previously (17,24,91). Briefly, HEK293 cells were maintained in 1x Dulbecco’s Modified Eagle Medium (Mediatech, Inc) without phenol red and supplemented with 10% Fetal Bovine Serum (FBS), 2mM L-Glutamine, and 1% penicillin/streptomycin (VWR, Calbiochem, and Invitrogen, respectively). Cells were transiently transfected with polyethyleneimine and RGS14-Luciferase(Luc) plus Gαi1-YFP (92) for 40 hrs. Cell were resuspended in Tyrode’s Solution (140 mM NaCl, 5 mM KCl, 1 mM MgCl_2_, 1 mM CaCl_2_, 0.37 mM NaH_2_PO_4_, 24 mM NaHCO_3_, 10 mM HEPES, and 0.1% glucose, pH 7.4) and plated at 10^5^ cells per well in a 96-well Optiplate (PerkinElmer Life Sciences). YFP expression was quantified using a TriStar LB 941 plate reader (Berthold Technologies) at 485 nm excitation and 530 nm emission. Next, 5 μM coelenterazine H (Nanolight Technologies) was incubated for 2 minutes and BRET measurements were taken at 485 nm (Luc emission) and 530 nm (YFP emission). The Net BRET ratio was quantified as follows: (530 nm signal / 485 nm signal) – (485 nm signal from Luc alone). Acceptor/Donor ratios were calculated as follows: (530 nm signal from YFP measurement)/(485 nm signal from BRET measurement). Each experiment was repeated 3 times.

### Dissociated hippocampal neuronal culture

Our protocol for culturing neurons was adapted from Beaudoin et al. 2012 (93). Brains were removed from E18-19 embryos obtained from a timed pregnant Sprague-Dawley rat (Charles River). The meninges were removed and the hippocampi were isolated in calcium and magnesium free HBSS (Invitrogen) supplemented with 1x sodium pyruvate (Invitrogen), 0.1 percent glucose (Sigma-Aldrich), and 10 mM HEPES (Sigma-Aldrich) pH 7.3. Isolated hippocampi were washed with the HBSS solution then dissociated using the same buffer containing 0.25 percent trypsin (Worthington) for 8 mins at 37°C. Trypsinized hippocampi were washed two times with the same HBSS buffer before being triturated 5-6 times with a fire-polished glass Pasteur pipette in BME (Invitrogen) supplemented with 10 percent FBS (VWR), 0.45 percent glucose (Sigma-Aldrich), 1x sodium pyruvate (Invitrogen), 1x Glutamax (Invitrogen), and 1x penicillin/streptomycin (HyClone). Neurons were counted and plated at a density of 80,000 cells/cm2 in the BME-based buffer on coverslips that had been etched with 70 percent nitric acid (Sigma-Aldrich) before being coated with 1 mg/mL poly-L-lysine (Sigma-Aldrich) in borate buffer. Cells were allowed to adhere for 1-3 hours before media was changed to Neurobasal (Invitrogen) supplemented with 1x B27 (Invitrogen) and 1x Glutamax (Invitrogen). Neurons were kept in a 37°C, 5 percent CO_2_ incubator and half of the media was replaced with new Neurobasal every 3-4 days until neurons were used for experiments.

### Confocal microscopy imaging

All reactions were carried out at room temperature. Cells were fixed with 4% paraformaldehyde for 10 minutes, then quenched with 0.75% glycine in 200 mM Tris pH 7.4, for 5 minutes. Cells were then permeabilized for 10 minutes in 0.1% Triton-X in PBS and stained with Hoechst (1:12,500 in PBS) for 4 minutes. Cells were finally washed three times in PBS and mounted onto slides with ProLong Diamond Antifade mountant (ThermoFisher). Images were taken on an Olympus FV1000 confocal microscope at 60x, then processed using ImageJ software. Approximately ten images per condition were obtained and representative images are shown.

### Hippocampal slice preparation and electrophysiological recordings

Hippocampal slices were prepared from wild type C57BL/6J mice (postnatal 5-8 days). Mouse pups were briefly placed in 70% ethanol, rapidly sacrificed by decapitation, and then brains rapidly removed. Recovered brains were sectioned into coronal slices (250 μm thick) by vibratome (Leica VT1200 S) in ice-cold cutting solution (MEM (GIBCO cat # 61100), NaHCO_3_ 26mM, HEPES 25mM, Tris 15mM, Glucose 10mM, MgCI_2_ 3mM). Under a dissecting microscope, hippocampal slices were removed and then carefully placed in a 0.4-μm pore size membrane insert (Transwell 3450-Clear, Corning Incorporated) 6-well plate. Slices (1-3 slices per well) were cultured at 37°C and 5% CO_2_ with the medium changed twice per week. Once cultured slices were stable, AAV-YFP-RGS14 (WT or mutant) was injected into CA1 and incubated for 10-14 days.

For electrophysiological recordings, slices were transferred to a recording chamber including ACSF containing (in mM): 124 NaCl, 2.5 KCl, 2 MgCl_2_, 2 CaCl_2_, 1.25 NaH2PO_4_, 26 NaHCO_3_ and 15 glucose. For the pairing induced LTP (pairing postsynaptic depolarization with presynaptic stimulation) experiments, the internal solution contained the following (in mM): 115 Cs-methanesulfonate, 20 CsCl, 2.5 MgCl_2_, 0.6 EGTA, 10 HEPES, 4 Na2-ATP, 0.4 Na2-GTP, and 10 phosphocreatine disodium salt. LTP was induced in whole-cell voltage clamp mode by pairing a post-synaptic depolarization to 0 mV with presynaptic stimulation delivered at 3 Hz (270 pulses over 1.5 minutes).

For imaging of slice cultures, slices were transferred to 4% paraformaldehyde (PFA) in phosphate buffered saline (PBS) for fixation at room temperature for 2 hrs, and then stored in PBS at 4°C until use. Twenty minutes prior to imaging, slices were incubated in 60% 2,2’-thiodiethanol to clear the tissue (94) and then imaged on a Zeiss 880 laser scanning confocal microscope (Carl Zeiss AG, Oberkochen, Germany). Data presented are pooled mean ± SD.

### Generation of RGS14 (L507R) CRISPR mouse line

To generate the Rgs14 point mutation, a CRISPR gRNA (CTGGTGGAGCTGCTGAATCGGG) targeting exon 15 along with a donor oligo (5’CAGCAGAATGGTCAAGCATTGGTGTAGGTGCTTTGGCCATATCGGCCCCTGACTCTG TGGCTCCCTCCAGGCCTGGTGGAGCTGC<u>G</u>G AATCG<u>A</u>GTGCAGAGCAGCGGGGCCCACGA TCAGAGAGGACTTCTTCGCAAAGAGGACCTGGTCCTTCCAGAATTTCTGCAGCTTCCTT CCCAA 3’; underlined bases are engineered) were designed to generate a CTG-to-CGG change creating the L507R (equivalent to L505R in human sequence and L504R in rat sequence) point mutation as well as G-to-A SNP conversion to create a PAM blocking mutation and G-to-A silent changes to create a novel TaqI restriction site in Rgs14 Mouse that could be used for screening.

To generate the RGS14 (L507R) mice, the gRNA (20 ng/ul), oligo donor (20 ng/ul), and CRISPR protein (20 ng/ul) were injected into 1-cell C57Bl/6J zygotes and subsequently transplanted at the 2-cell stage into C57Bl/6J pseudopregnant females by the Emory Mouse Transgenic and Gene Targeting Core. Genomic DNA from toes was amplified via PCR using primers: FW 5’-GTGGCATCAGAGAGGCCTG-3’ and RV 5’-GTTACACAGATGCCAGAGGAC-3’. PCR bands at 387bp (wild type), 188bp, and/or 199 (mutant) were produced. Of the 95 pups born, 9 of them (approx. 10%) were positive for the mutation. Two positive animals were selected initially for continued breeding. To dilute out/eliminate potential off-target mutations due to CRISPR/Cas9 gene editing, each animal was back-crossed (Het-Het) against a fresh wild type C57Bl/6J background for three generations, and then a single line was continued for back-crossing for five generations prior to behavioral tests.

### Morris water maze behavioral test

Morris Water Maze was conducted as described previously (20). Adult wild type RGS14 and homozygous RGS14 LR mice (littermates ages 2–6 mo) were used. The water maze consisted of a circular swim arena (diameter of 116 cm, height of 75 cm) surrounded by extramaze visual cues that remained in the same position for the duration of training and filled to cover the platform by 1cm at 22°C. Water was made opaque with nontoxic, white tempera paint. The escape platform was a circular, nonskid surface (area 127 cm^2^) placed in the NW quadrant of the maze. Acquisition training consisted of 5 test days with four daily trials. Mice entered the maze facing the wall and began each trial at a different entry point in a semirandom order. Trials lasted 60 s or until the animal mounted the platform with a 15-min intertrial interval. A probe trial was conducted on day 6 wherein the platform was removed, and the animal swam for 60 s and the time spent in the target quadrant (NW) versus the adjacent and opposite quadrants was recorded. A video camera mounted above the swim arena and linked to TopScan software recorded swim distance, swim speed and time to platform and was used for tracking and analysis. Statistics were ANOVA and post hoc Dunnett’s test.

### RGS14 LR CRISPR mouse immunofluorescence

Male RGS14 WT (n=3) and RGS14 LR (n=3) mice were euthanized with an overdose of sodium pentobarbital (Fatal Plus, 150 mg/kg, i.p.; Med-Vet International, Mettawa, IL) and transcardially perfused with 4% paraformaldehyde dissolved in 0.01 M PBS. After extraction, brains were postfixed for 24 h in 4% paraformaldehyde at 4°C and subsequently transferred to 30% sucrose/PBS solution for 72 h at 4°C. Brains were embedded in OCT medium (Tissue-Tek; Sakura, Torrance, CA) and serially sectioned by cryostat (Leica) into 40-µm coronal slices. Brain sections were stored in 0.01 M PBS (0.02% sodium azide) at 4°C before IHC.

Immunohistochemistry (IHC) was performed to visualize RGS14 protein in brain sections at the level of the hippocampus and striatum. Brain sections from mice of both genotypes were subjected to 3 min antigen retrieval at 100°C in 10 mM sodium citrate buffer. Following antigen retrieval, brain sections were blocked for 1 h at room temperature in 5% normal goat serum (NGS; Vector Laboratories, Burlingame, CA) diluted in 0.01 M PBS/0.1% Triton-X permeabilization buffer. Sections were then incubated for 24 h at 4°C in this same NGS blocking/permeabilization buffer now including a mouse primary antibody raised against RGS14 (NeuroMab; clone N133/21) at 1:500 dilution. After washing, sections were then incubated for 2 h in blocking/permeabilization buffer with goat anti-mouse AlexaFluor 568 (Invitrogen, Carlsbad, CA) at 1:500 dilution. After washing, the sections were mounted onto Superfrost Plus slides (Thermo Fisher Scientific, Norcross, GA) and coverslipped with Fluoromount-G plus DAPI (Southern Biotech, Birmingham, AL).

Fluorescent micrographs of immunostained sections were acquired on a Leica DM6000B epifluorescent upright microscope at 5, 10, or 20x magnification. Micrographs were acquired for dorsal hippocampal Cornu Ammonis fields CA1 and CA2, piriform cortex (Pir Ctx), central amygdala (CeA), dorsal striatum (DS), and nucleus accumbens (NAc). For each regional comparison of RGS14 immunostaining between genotypes, uniform exposure parameters were used. All images were processed with ImageJ (NIH) to reduce background and enhance contrast.

### Next generation RNA-sequencing and data analysis

Cultured rat hippocampal neurons (DIV18) were infected with AAV2/9 virus expressing either RGS14 WT (n=4), RGS14 carrying mutations RKK/AAA (NLSm) (n=3) or RGS14 LL/AA (NESm) (n=3). Infected neurons then were harvested and RNA collected (Qiagen RNeasy Mini Kit). Purified RNA (500 ng) were converted to sequencing libraries using the KAPA mRNA HyperPrep kit. To perform RNA-seq, libraries were pooled at equimolar ratio and sequenced on a NextSeq500 using 75 bp paired-end sequencing. Data processing involved raw fastq reads were mapped to the rn6 version of the rat genome. High percentage of reads mapped (> 90%) and were unique (orange bar), indicating high quality libraries. Fastq files were mapped to the RGSC 6.0/rn6 genome using STAR (95) and gene coverage for all exons annotated using GenomicRanges (96). To identify detected transcripts genes are filtered for detection based on being expressed at > 3 reads per million in at least 3 samples (first blue dotted line) resulting in the detection of 10,324 out of 17,324 transcripts. To analyze samples for differential gene expression and sample variation, all detected genes were used as input for edgeR v3.24.3 (97) to identify differentially expressed genes. Data sets were subjected to principal component analysis of variation using the princomp function in R v3.5.2 between samples to examine interclustering and differences between the groups.

## Acknowledgements

The authors would like to acknowledge and thank Dr. YuhMin Chook for generously providing the CRM1 (XPO1) cDNA construct and guidance on its interaction conditions. We would also like to thank Dr. Jeremy Boss for his constructive input and contributions regarding RNA sequencing.

