## Supporting Information for "Human genetic variants disrupt RGS14 nuclear shuttling and regulation of LTP in hippocampal neurons"

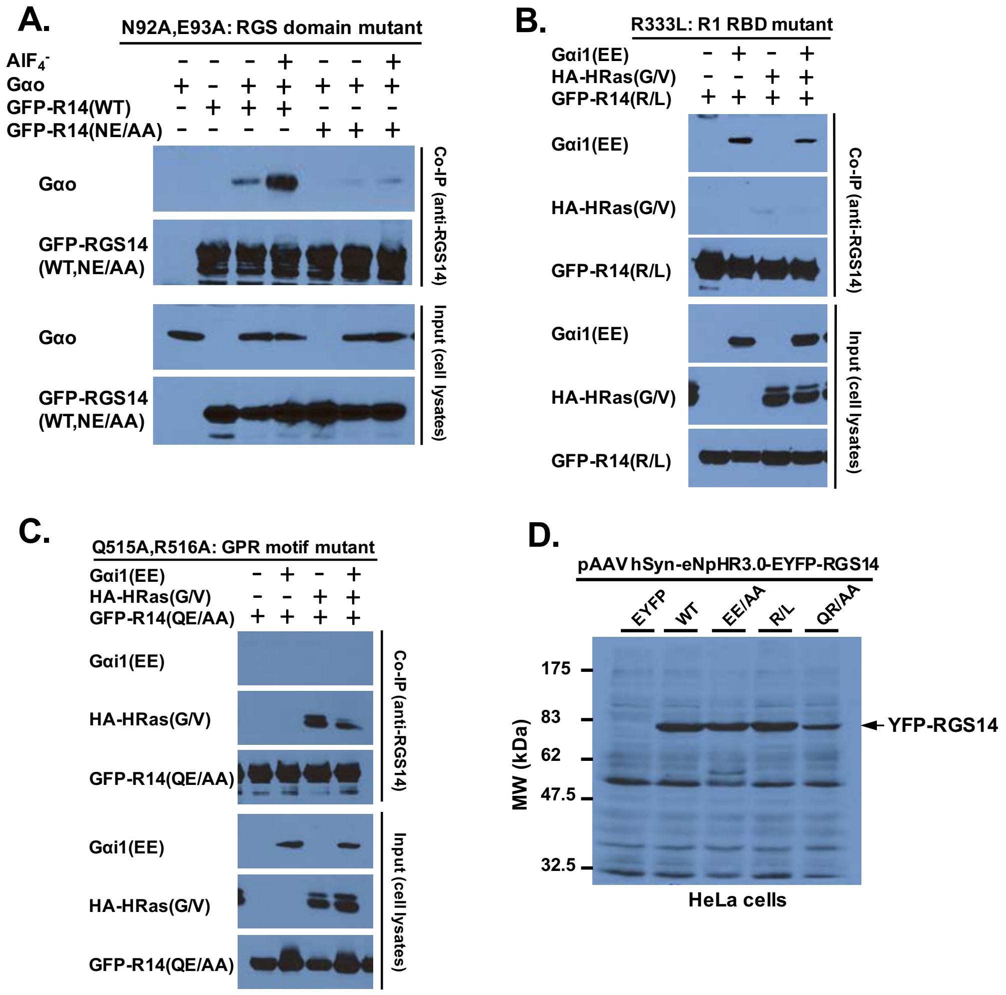


**Figure S1. Functional characterization of targeted loss-of-function mutations in RGS14 and generation of wild type and mutant RGS14 Adeno-associated viral (AAV) vectors.**

***A)*** *The RGS domain loss-of-function mutant of RGS14. Amino acids N92 and E93 in the rat RGS14 sequence N-terminally fused to GFP were both mutated to alanine (NE/AA). HeLa cells were transfected with either wild type (WT) or mutant (NE/AA) GFP-RGS14 alone or co-transfected together with Gαo, and then tested for functional RGS14:Gαo binding in the absence or presence of aluminum fluoride (AlF4-). RGS14 was immunoprecipitated (IP) from cell lysates and the recovered samples subjected to SDS-PAGE and immunoblotted for the presence of RGS14 (anti-GFP) or Gαo (anti-Gαo).* ***B)*** *The R1 Ras-binding domain (RBD) loss-of-function mutant of RGS14. Amino acid R333 in the rat RGS14 sequence N-terminally fused to GFP was mutated to leucine (R/L). HeLa cells were transfected with either wild type (WT) or mutant (R/L) GFP-RGS14 alone or co-transfected together with either Glu-Glu epitope tagged Gαi1 (Gαi1-EE) or constitutively activated HA-tagged H-Ras-G12V (HA-HRas(G/V)), and then tested for functional RGS14 binding. RGS14 was immunoprecipitated (IP) from cell lysates and the recovered samples subjected to SDS-PAGE and immunoblotted for the presence of RGS14 (anti-GFP) and either Gαi1 (anti-EE) or H-Ras(GV) (anti-HA).* ***C)*** *The GPR motif loss-of-function mutant of RGS14. Amino acids Q515 and R516 in the rat RGS14 sequence N-terminally fused to GFP were both mutated to alanine (QE/AA). HeLa cells were transfected with either wild type (WT) or mutant (QE/AA) GFP-RGS14 alone or co-transfected together with either Gαi1-EE or Glu-Glu epitope tagged Gαi1 (Gαi1-EE) or constitutively* *activated HA-tagged H-Ras-G12V(HA-HRas(G/V)), and then tested for functional RGS14 binding. RGS14 was immunoprecipitated (IP) from cell lysates and the recovered samples subjected to SDS-PAGE and immunoblotted for the presence of RGS14 (anti-GFP) and either Gαi1 (anti-EE) or H-Ras(GV) (anti-HA).* ***D)*** *Wild type RGS14 and each of the RGS14 mutants NE/AA, R/L and QE/AA described above (A-C) were cloned into the adeno-associated viral vector pAAV hSyn-eNpHR3.0-EYFP. RGS14 and mutants were placed in frame with eYFP. Empty vector (eYFP), or vector expressing RGS14 (WT), or RGS14 mutants EE/AA, R/L and QR/AA were each transfected into HeLa cells to confirm protein expression of the correct size. eYFP-RGS14 was visualized as a single band of the correct size. Due to low/inefficient expression of proteins in HeLa cells, the immunoblot was overexposed to allow visualization of RGS14 bands, increasing non-specific staining due to secondary antibody. These cDNA vectors were used to generate AAV2/9-YFP-RGS14 wild type and mutant viruses for subsequent studies as described.*

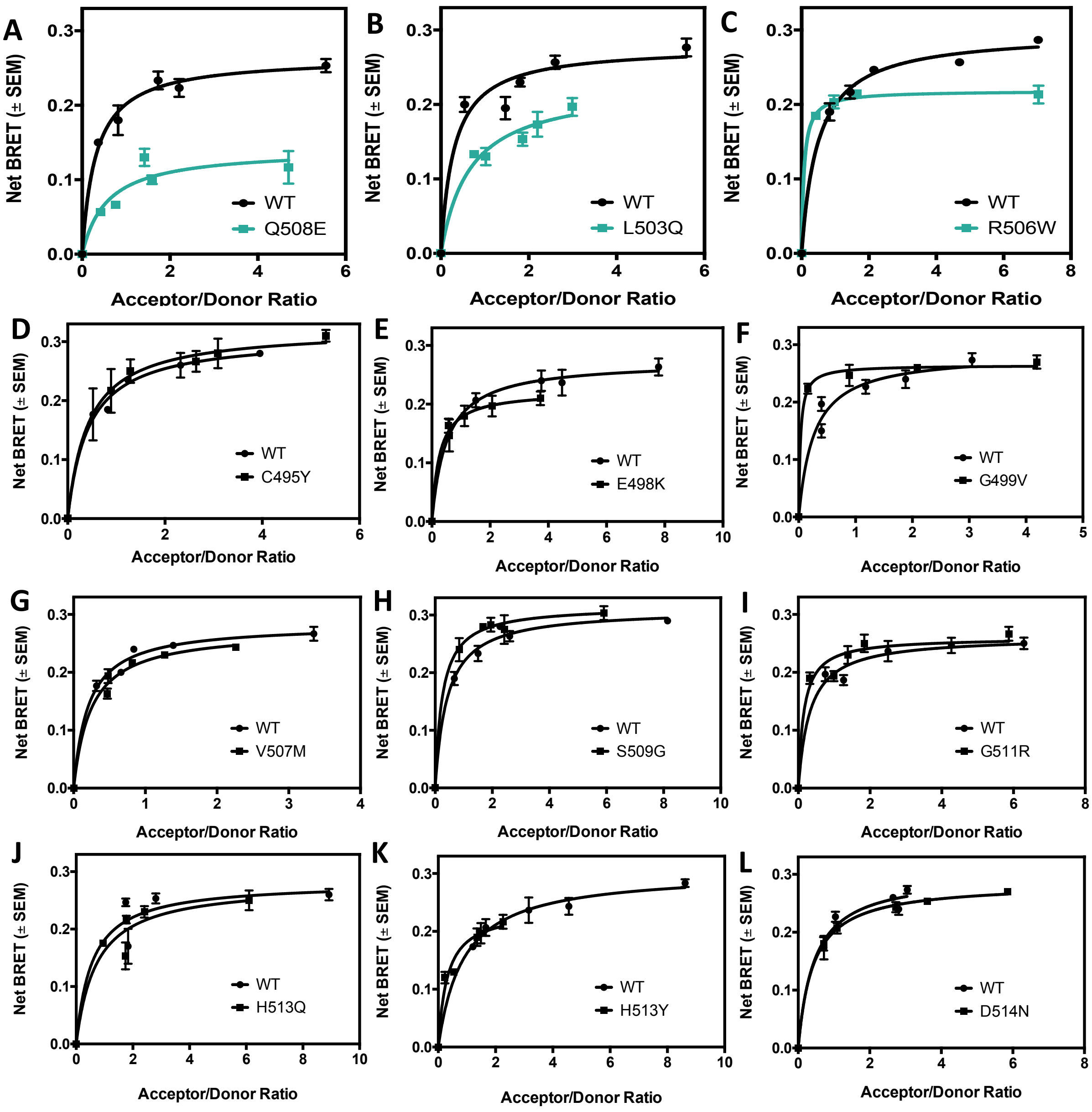


**Figure S2. Most human variants in RGS14 do not affect association with Gαi1-GDP.**
*Variants from Figure 2 were tested for their capacity to associate with Gαi1-GDP, as measured by Net BRET. There were several variants that exhibited a reduction in Gαi1-GDP association* ***(A-C shown in teal)****, but others had no robust phenotype* ***(D-L)****. All experiments were n=3 and expressed as mean +/-SEM.*

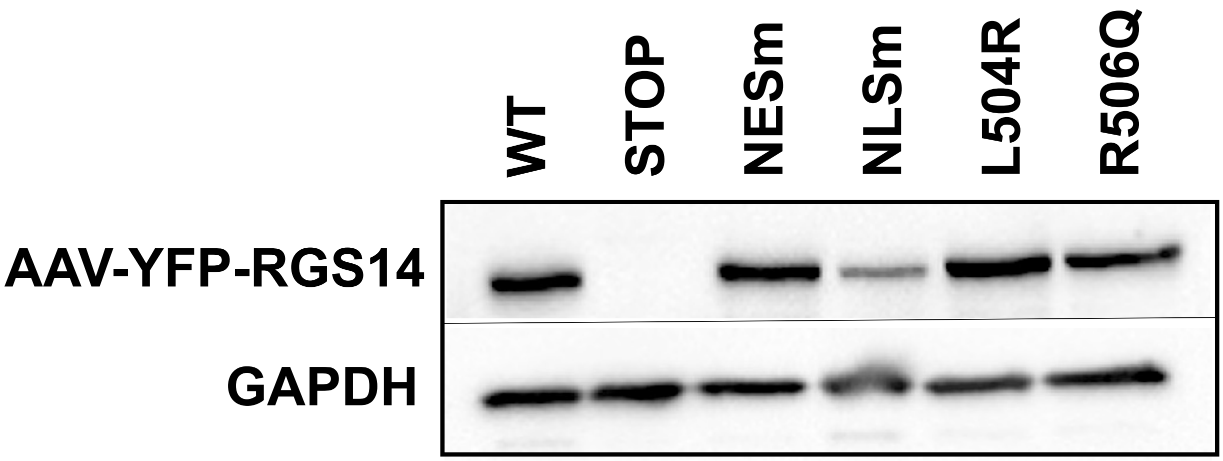


**Figure S3. Expression of RGS14 constructs is verified by immunoblot.**
*Wild type RGS14 expressed a single band correlating to the approximate size of YFP-RGS14 (~85 kDa), while the STOP mutation correlating to YFP-STOP-RGS14 produced no detectable RGS14 as expected. The NES, NLS, RQ, and LR mutations similarly produced a single band corresponding to full length YFP-RGS14. GAPDH was used as a loading control.*

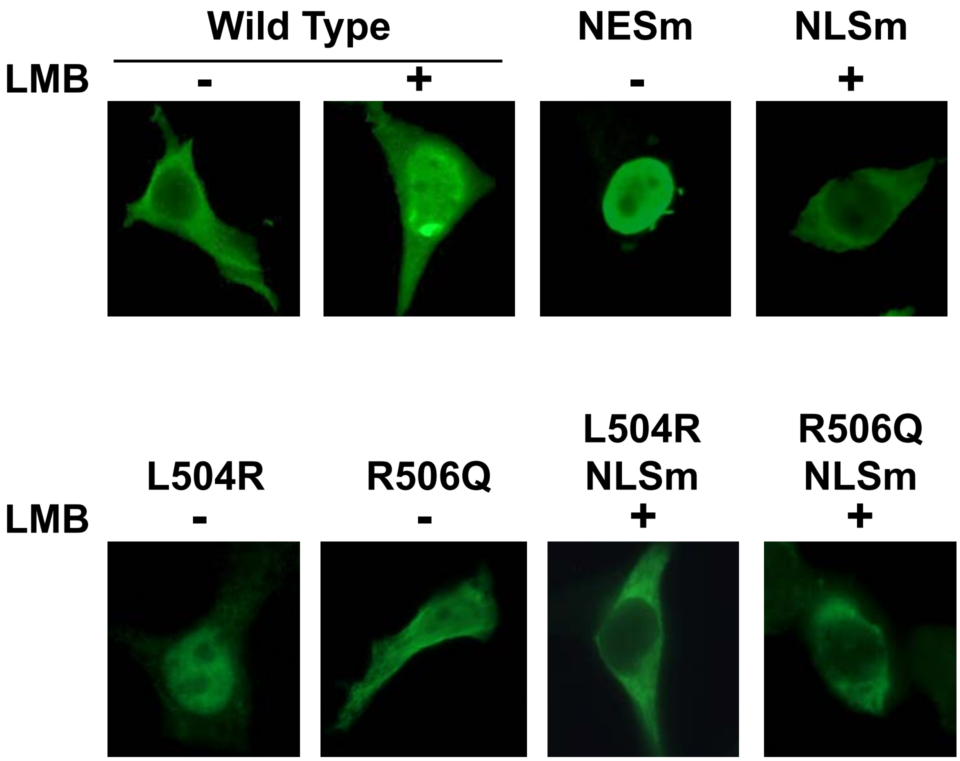


**Figure S4. Nuclear localization mutations control nuclear trafficking and/or export of RGS14 in hippocampal neurons.**

*Wild type RGS14 translocates to the nucleus of hippocampal neurons following LMB treatment (top left). RGS14 NESm is sequestered in the nucleus under vehicle conditions, and NLSm fails to translocate to the nucleus, even in the presence of LMB (top right). While L504R and R506Q have nuclear localization phenotypes to varying degrees (bottom left), adding an NLS mutation to each construct entirely prevents nuclear localization (bottom right).*

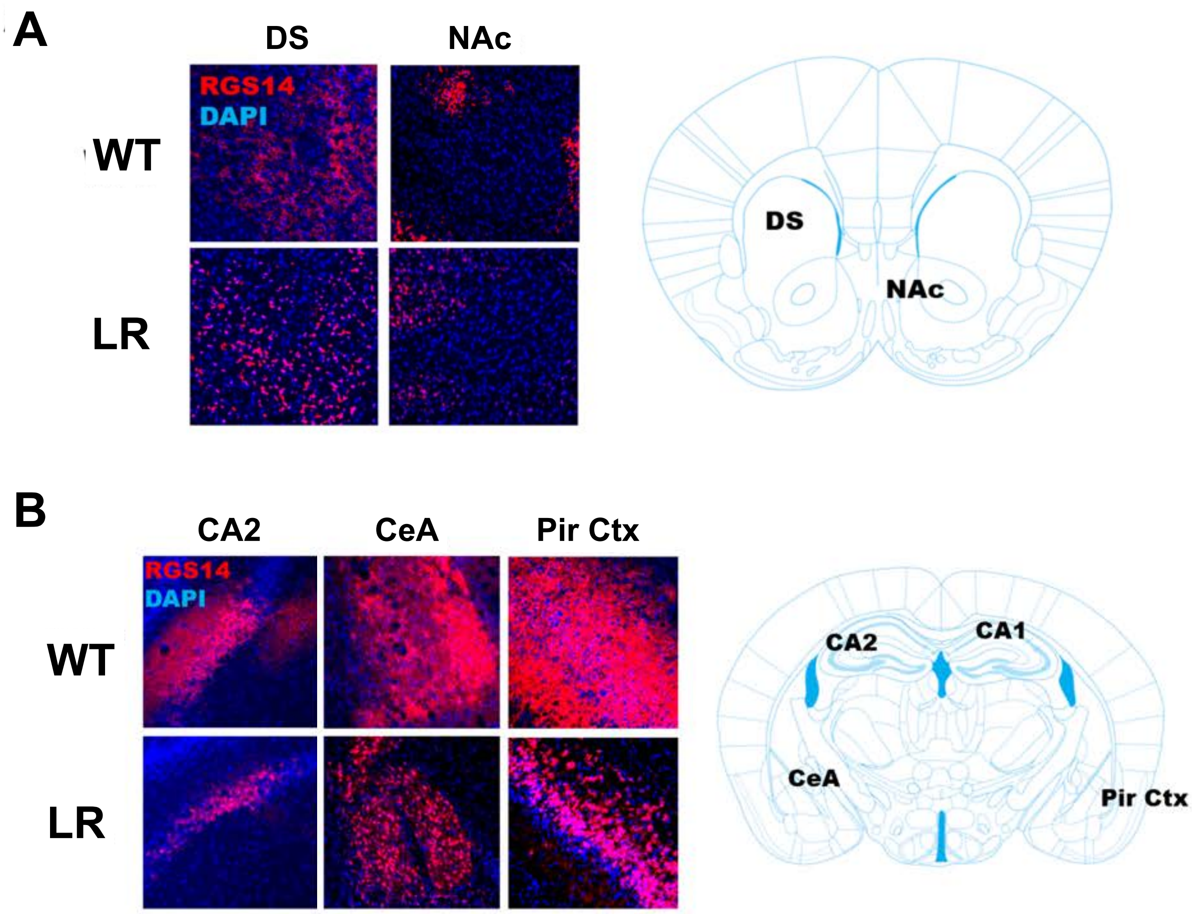


**Figure S5. RGS14 LR mice unveil RGS14 expression outside of the hippocampus.**
***A)*** *RGS14 expression in dorsal striatum (DS) and nucleus accumbens (NAc) is shown in red. DAPI confirms nuclear sequestration of RGS14 LR by immunofluorescence.* ***B)*** *Distinct nuclear localization facilitated RGS14 identification outside of the hippocampus, unveiling more widespread expression than previously appreciated, including the central amygdala (CeA) and piriform cortex (Pir Ctx). WT expression was confirmed for these regions, indicating this was not an artefact of the mutation itself. Images are representative of 3 WT mice and 3 LR mice.*

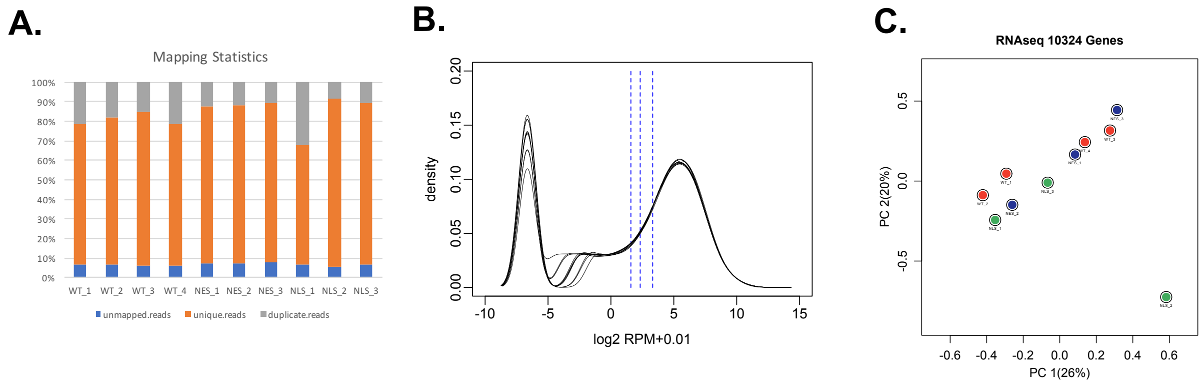


**Figure S6. Expression of RGS14 or RGS14 mutants excluded from the nucleus (NLSm) or trapped in the nucleus (NESm) do not alter mRNA transcripts in cultured hippocampal neurons at rest.**
*Cultured rat hippocampal neurons (DIV18) were infected with AAV2/9 virus expressing either RGS14 WT (n=4), RGS14 carrying mutations RKK/AAA (NLSm) (n=3) or RGS14 LL/AA (NESm) (n=3) for 2 weeks. Protein expression was observed as shown in Figure S4.* ***A)*** *Raw FASTq sequencing reads plotted as percentages of total. Unique reads are presented as orange.* ***B)*** *Identification of detected transcripts. Genes were filtered based on being expressed at >3 reads per million in at least 3 samples (blue dotted lines). 10,324 of ~17,000 transcripts were detected.* ***C)*** *Differential analysis and Sample Variation. No differentially expressed genes were detected in any comparison (WT=Orange, NLSm=Green, and NESm=Blue). Principle component analysis of variation between samples shows they intercluster with no differences between the three groups.*

| Variant | Forward | Reverse |
|---|---|---|
| **Q508E** | ctgaatcgtgtggagagcagcggggcc | ggccccgctgctctccacacgattcag |
| **L503Q** | gaaggcctagtggagcagctgaatcgtgtgcag | ctgcacacgattcagctgctccactaggccttc |
| **R506W** | gtggagctgctgaattgggtgcagagcagc gggg | cccc gctgctctgc acccaattca gcagctccac |
| **C495Y** | gaaa cgacag acc tacgacattgaag | cttcaatgtcgtaggtctgtcgtttc |
| **E498K** | cagacctgcgacattaaaggcctagtggagctgctg | cagcagctccactaggcctttaatgtcgcaggtctg |
| **G499V** | cgacagacctgcgacattgaagtcctagtggagctgctgaatcgt gtgc | gcacacgattcagcagctccactaggacttcaatgtcgcaggtct gtcg |
| **V507M** | gctgaatcgtatgcagagcagcg | cgctgctctgcatacgattcagc |
| **S509G** | gaatcgtgtgcagggcagcggggcccatg | catgggccccgctgccctgcacacgattc |
| **G511R** | cgtgtgcagagcagcagggcccatgaccagagg | cctctggtcatgggccctgctgctctgcacacg |
| **H513Q** | agcagcggggcccaggaccagaggggac | gtcccctctggtcctgggccccgctgct |
| **H513Y** | agcagcggggcctatgaccagaggggac | gtcccctctggtcataggccccgctgct |
| **D514N** | ccaaagctgaagcctggaaaggacctgccgctcggtgtggaag ag | ctcttccacaccgagcggcaggtcctttccaggcttcagctttgg |
| **R506Q** | ggcctagtggagctgctgaatcaggtgcagagcagcggggccc atgacc | ggtcatgggccccgctgctctgcacctgattcagcagctccacta ggcc |
| **L504R** | gaaggcctagtggagctgcggaatcgtgtgcagagcagc | gctgctctgcacacgattccgcagctccactaggccttc |
| **NESm (L503A/ L504A)** | gaa ggc cta gtg gag gcg gcg aat cgt gtg cag agc | gct ctg cac acg att cgc cgc ctc cac tag gcc ttc |
| **NLSm (R208A/ K209A/ K210A)** | ccg gac act gcg gcg gcg gcg cca aag ctg aag | ctt cag ctt tgg cgc cgc cgc cgc agt gtc cgg |
| **GPRm (Q515A/ R516A)** | ggggcccatgacgccggccggacttcttcgcaaag | ggggcccatgacgccggccggacttcttcgcaaag |
| **RGS-null (E92A/ N93A)** | aaggaattcagcgccgccgccgtaactttctggcaagc | aaggaattcagcgccgccgccgtaactttctggcaagc |
| **R1-null (R333L)** | ggc atc tgt gag aag cta ggc ctc tct cta cct g | ggc atc tgt gag aag cta ggc ctc tct cta cct g |

**Table S1. Primers used for RGS14 Human Variants and Functional Mutations.**
